# Evolutionary rate covariation is pervasive between glycosylation pathways and points to potential disease modifiers

**DOI:** 10.1101/2024.05.03.592434

**Authors:** Holly J. Thorpe, Raghavendran Partha, Jordan Little, Nathan L. Clark, Clement Y. Chow

## Abstract

Mutations in glycosylation pathways, such as N-linked glycosylation, O-linked glycosylation, and GPI anchor synthesis, lead to Congenital Disorders of Glycosylation (CDG). CDGs typically present with seizures, hypotonia, and developmental delay but display large clinical variability with symptoms affecting every system in the body. This variability suggests modifier genes might influence the phenotypes. Because of the similar physiology and clinical symptoms, there are likely common genetic modifiers between CDGs. Here, we use evolution as a tool to identify common modifiers between CDG and glycosylation genes. Protein glycosylation is evolutionarily conserved from yeast to mammals. Evolutionary rate covariation (ERC) identifies proteins with similar evolutionary rates that indicate shared biological functions and pathways. Using ERC, we identified strong evolutionary rate signatures between proteins in the same and different glycosylation pathways. Genome-wide analysis of proteins showing significant ERC with GPI anchor synthesis proteins revealed strong signatures with ncRNA modification proteins and DNA repair proteins. We also identified strong patterns of ERC based on cellular sub-localization of the GPI anchor synthesis enzymes. Functional testing of the highest scoring candidates validated genetic interactions and identified novel genetic modifiers of CDG genes. ERC analysis of disease genes and biological pathways allows for rapid prioritization of potential genetic modifiers, which can provide a better understanding of disease pathophysiology and novel therapeutic targets.

**AUTHOR SUMMARY:** Congenital Disorders of Glycosylation (CDGs) are a group of rare disorders resulting from impaired protein glycosylation. Glycosylation is the addition of sugar chains onto proteins and is required for proper protein function. CDG patients typically present with seizures and hypotonia. However, they can have a large amount of clinical variability, which is likely influenced by modifier genes. Modifier genes are genes that affect a phenotype without causing the disease. Using an evolutionary method that examines proteins that evolve at similar rates, we identified proteins within glycosylation pathways and among other unexpected pathways, such as ncRNA modification and DNA repair, that could be potential genetic modifiers of CDG genes. We also tested top protein pairs using the *Drosophila* eye as a model and identified novel genetic modifiers of CDG genes. Broadening our understanding of CDG modifiers can help us to better understand why loss of glycosylation results in specific patient symptoms and could provide new treatment targets.

## INTRODUCTION

Phenotypic variation is common in Mendelian diseases, even in patients with the same disease-causing variant (1–3). Understanding the factors underlying variable disease presentation can be difficult despite often knowing the causal disease gene. Genetic modifiers can enhance or suppress the phenotype of a primary disease-causing mutation, but they may not independently lead to disease phenotypes.

Congenital disorders of glycosylation (CDG) are a group of ultra-rare multisystemic disorders typically characterized by seizures, hypotonia, and neurodevelopmental delays (4,5). However, because of the ubiquity of glycosylation, symptoms of CDGs can affect most organ systems. Loss-of-function mutations in >150 glycosylation genes lead to CDGs (6). Patients with the same CDG and even the same disease-causing variant can show phenotypic variability, indicating that genetic background and environment can influence clinical presentation (7,8). Current treatment options for CDGs are limited, primarily focusing on symptom management (9,10). Identifying genetic modifiers of glycosylation genes could lead to new therapeutic targets for CDG patients.

Protein glycosylation is essential for proper protein folding, stability, and localization (4,5,11). There are three main types of glycosylation: N-linked glycosylation, O-linked glycosylation, and glycosylphosphatidylinositol (GPI) anchor biosynthesis. N-linked glycosylation is the addition of a glycan onto an asparagine (4,5). The glycan is synthesized on the endoplasmic reticulum (ER) membrane, with the stepwise addition of sugars onto a dolichol-phosphate. The glycan is then transferred from the dolichol-phosphate onto a protein and undergoes further glycan additions and trimming in the ER and Golgi. O-linked glycosylation is the addition of a glycan to the hydroxyl group of a serine, threonine, or hydroxylysine (4). O-linked glycosylation encompasses the most extensive variety of glycans and can occur in the ER, Golgi, or cytoplasm (12). O-linked glycans are classified by the first sugar attached to the protein. The rest of the glycan is built directly onto that sugar.

GPI anchors are glycolipids that tether proteins to the cell membrane (8,13). GPI anchor synthesis begins by adding an N-acetylglucosamine (GlcNAc) onto a phosphatidylinositol on the cytoplasmic side of the ER membrane. The acyl group is then removed before the glycan is flipped onto the lumen side of the ER membrane. In the ER lumen, three mannoses (four in some species/tissues) and three phosphoethanolamine groups are added before a protein is attached to the GPI anchor. The GPI anchor then undergoes fatty acid remodeling before it is sent to the Golgi for further remodeling. Then, the GPI-anchored glycoprotein is trafficked to the cell surface.

Standard methods for identifying genetic modifiers typically involve utilizing natural variation in the population or performing large mutagenesis screens (1). Several recent studies reported modifier screens in animal models of specific CDGs (14–16). However, with over 150 CDGs, performing similar studies for each CDG is impractical. PMM2-CDG, the most common CDG, has just over 1000 diagnosed patients worldwide (17). Most CDGs, however, have fewer than 100 patients, making it challenging to have enough statistical power to distinguish impactful variation within the population (8,18). It is likely CDGs share common genetic modifiers because different forms can have similar effects on glycosylation and similar clinical presentations. Instead of examining each CDG independently, we sought to identify common modifier genes for all or a subset of CDGs. To do this, we used evolutionary rate covariation (ERC), a computational evolutionary method developed to identify proteins with evolutionary rates that covary across species (19). Protein pairs with high ERC values have similar changes in evolutionary rate and are thought to function together, either in a complex or in a pathway, and may genetically modify each other. ERC allows us to identify proteins with evolutionary rates that covary with CDG proteins, potentially identifying genetic modifiers.

A number of recent studies suggest that other glycosylation genes modify CDG genes. In 2002, a study examined a small group of PMM2-CDG patients, the most common CDG, and identified a variant in *ALG6*, an N-linked glycosylation gene, segregating with more severe outcomes (20). A modifier screen and ERC analysis of NGLY1, a deglycosylating enzyme associated with a CDG, identified multiple glycosylation genes showing high ERC with NGLY1, implicating them as potential modifiers (15). A CRISPR screen in cells with reduced DPAGT1 function, the first step in N-glycan synthesis, identified multiple glycosylation genes able to rescue glycosylation defects caused by inhibition of DPAGT1 (14). In yet another study, a yeast model of PMM2-CDG was evolved over 1000 generations to identify genetic changes that increased the viability of the *PMM2* mutant yeast strains (16). Several top genes that accumulated compensatory mutations were also involved in glycosylation. These studies indicate that genetic variation in glycosylation genes might potentially modify the outcomes of CDGs.

Applying a computational evolutionary method allows for more rapid and broad detection of disease modifiers. Identifying genetic modifiers of CDGs will lead to a better understanding of the physiology underlying variable symptoms in patients and could point to new therapeutic targets. Using ERC, we performed an analysis to discover proteins that have correlated evolutionary rates with glycosylation proteins to discover genetic modifiers of CDGs. We found strong intra-pathway ERC signatures among glycosylation proteins in each of the three glycosylation pathways, with the highest average score in GPI anchor synthesis. There were also high inter-pathway ERC values between proteins in different glycosylation pathways. Strikingly, GPI anchor synthesis proteins had many high ERC values with N-linked glycosylation proteins. Protein complexes and proteins with similar functions in glycosylation were also enriched for strong ERC. When examining the genome for non-glycosylation modifiers of GPI anchor synthesis proteins, we found high ERC scores with proteins involved in multiple unrelated pathways, including ncRNA and DNA repair. We found that the strength of these ERC scores is driven by the GPI anchor synthesis enzyme’s location in the cell. *In vivo* testing of top scoring protein pairs, including GPAA1 with ALG1 and PIGA with RBSN, showed a genetic interaction in a *Drosophila* eye model. Using computational evolutionary methods, we identified patterns of ERC between CDG proteins and novel potential modifier proteins, many of which validated *in vivo*.

## RESULTS

### High ERC is pervasive between glycosylation proteins

To find potential candidate genetic modifiers of CDGs, we created a dataset of ERC scores across the genome. Using evolutionary rate data from yeast, nematode worms, *Drosophila*, vertebrates, and mammals (S1 Fig A), we calculated ERC scores for 19,149 proteins across the genome resulting in 180,112,645 pairwise ERC values (S1 Fig B). The genome-wide dataset values range from -10.73 – 19.30, with an average of 0.12 ± 1.47. Higher values indicate stronger covariation of evolutionary rates. In this study, ERC values ≥ 3 are considered notably elevated because they are two standard deviations from the mean. Using this dataset, we compared ERC scores between subsets of proteins.

We first examined the pairwise correlated evolutionary rates among all glycosylation proteins. We included 296 glycosylation proteins in our analyses (43,660 protein pairs), including all known CDG proteins and glycosylation proteins not yet associated with a disease (S1 Data). These include enzymes and support proteins involved in synthesizing and modifying glycans, enzymes required to form glycosylation building blocks, and proteins involved in transporting glycoproteins throughout the cell. We calculated ERC scores between all 296 glycosylation proteins (S2 Fig, S1 Data). Glycosylation protein pairs showed higher average values of ERC than expected by chance (mean ERC = 0.41, p < 1×10^-5^ Permutation test) (S3 Fig, Table 1). The mean ERC score for glycosylation protein pairs is more than 12 standard deviations above the mean of the permutations.

**Table 1:** Average pairwise ERC scores in glycosylation pathway.

| Table 1: Average pairwise ERC scores in glycosylation pathway |  |  |  |  |
| --- | --- | --- | --- | --- |
| Pathway | Proteins in pathway | Pairwise scores | Average ERC score | P-value |
| All Glycosylation | 296 | 43,660 | 0.41 | $<1 \times 10^{-5}$ |
| GPI Anchor Synthesis | 27 | 351 | 1.69 | $<1 \times 10^{-5}$ |
| N-linked Glycosylation | 71 | 2,485 | 0.64 | $<1 \times 10^{-5}$ |
| O-linked Glycosylation | 122 | 7,381 | 0.31 | $<1 \times 10^{-5}$ |
| Other | 76 | 2,850 | 0.47 | $<1 \times 10^{-5}$ |

Average of the pairwise ERC scores between the proteins in each pathway. Significance tested against 100,000 permutations.

Proteins in a pathway are often under similar evolutionary constraints and typically have high ERC values (19). To determine if each of the individual glycosylation pathways exhibited even higher ERC than when all glycosylation proteins are considered, we analyzed pairwise ERC of proteins in each major glycosylation pathway (GPI anchor synthesis, N-linked glycosylation, and O-linked glycosylation) as well as proteins that contribute to multiple glycosylation pathways (Other) (Table 1). As before, permutation testing was used to determine the significance of the ERC values. GPI anchor synthesis had a mean ERC score of 1.69 (27 proteins, p < 1×10^-5^) (Figure 1), N-linked glycosylation had a mean ERC score of 0.64 (71 proteins, p < 1×10^-5^) (S4 Fig A), and O-linked glycosylation had a mean ERC score of 0.31 (122 proteins, p < 1×10^-5^) (S4 Fig B). The proteins involved in multiple pathways (Other) also showed higher than expected ERC with an average of 0.47 (76 proteins, p < 1×10^-5^) (S4 Fig C). Strikingly, the average score of the GPI anchor synthesis pathway was nearly four times as high as the average of all glycosylation proteins (GPI average = 1.69, all glycosylation average = 0.41). O-linked glycosylation (average ERC = 0.31) is the only pathway with a lower average ERC score than the overall glycosylation average. This lower score is likely because O-linked glycosylation comprises eleven independent pathways that form different types of O-linked glycans, reducing the number of proteins in this category that work directly with each other.

**Figure 1:**
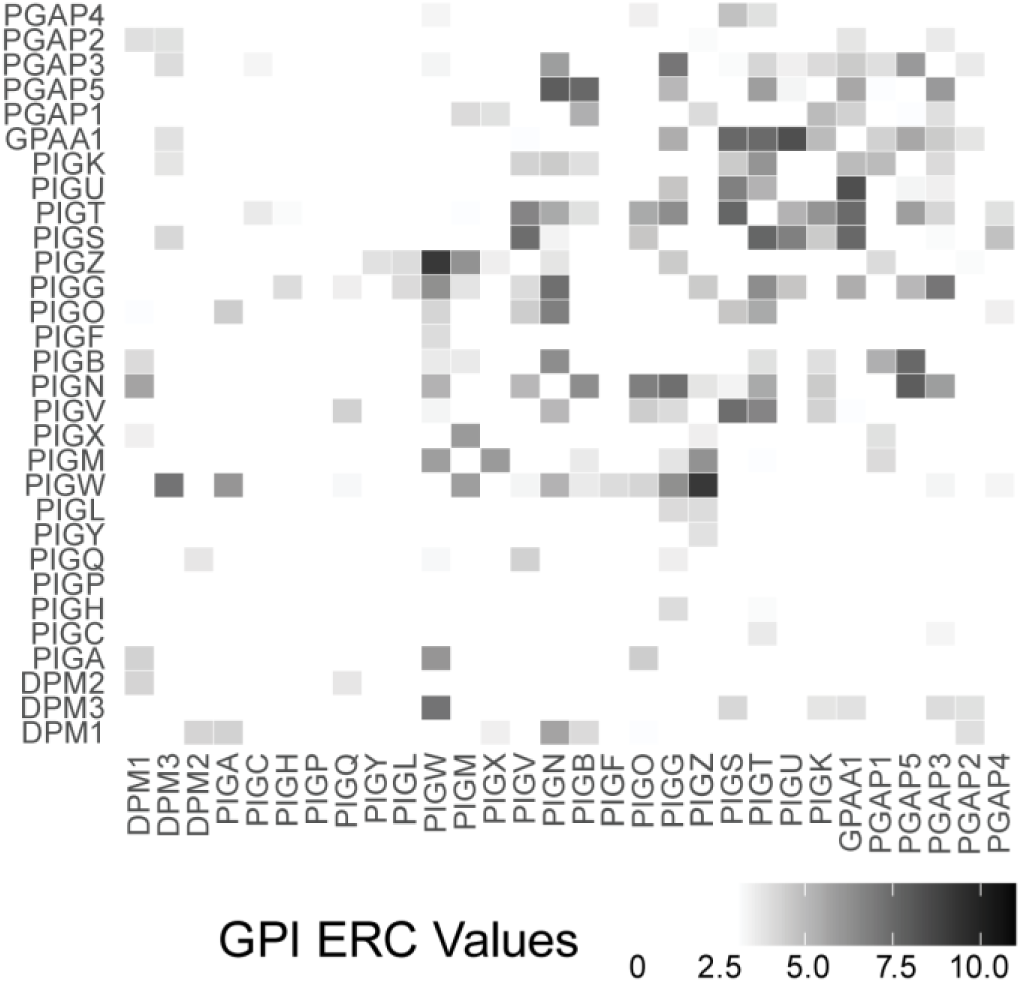
Heatmap showing significant (>3) pairwise ERC scores between proteins involved in GPI anchor synthesis.

### Strong evolutionary covariation between proteins in different glycosylation pathways

Glycosylation is a broad term encompassing multiple pathways with hundreds of proteins in different organelles. While some proteins associated with CDGs impact different types of glycans, nearly all enzymatic proteins are specific to one of the three main glycosylation pathways. We asked if the high mean ERC score observed across all glycosylation proteins (mean ERC = 0.41, p < 1×10^-5^) was driven primarily by high scores between proteins in the same pathway or if high ERC scores between proteins in different pathways were also observed. Proteins in all three glycosylation pathways showed strong ERC with proteins in different pathways (Table 2, S5 Fig, S6 Fig). As within the pathway, GPI anchor synthesis had the highest scores with other glycosylation pathways. The average ERC score between proteins in GPI anchor synthesis and proteins in N-linked glycosylation is 0.81 (p < 1×10^-5^), O-linked glycosylation is 0.46 (p < 0.014), and Other is 0.61 (p < 1.30^-4^). The average scores between GPI anchor synthesis and either N-linked, O-linked glycosylation, or Other are higher than the average of all glycosylation proteins (mean ERC = 0.41). This strong signal between glycosylation proteins in different pathways, especially those in GPI anchor synthesis and N-linked glycosylation, could indicate unexpected interactions between pathways.

**Table 2:** Average ERC values between glycosylation pathways.

| Table 2: Average ERC values between glycosylation pathways |  |  |  |  |
| --- | --- | --- | --- | --- |
|  | GPI | N-linked | O-linked | Other |
| GPI | 1.69<br>$p < 1 \times 10^{-5}$ | - | - | - |
| N-linked | 0.81<br>$p < 1 \times 10^{-5}$ | 0.64<br>$p < 1 \times 10^{-5}$ | - | - |
| O-linked | 0.46<br>$p < 1.44 \times 10^{-2}$ | 0.42<br>$p < 1 \times 10^{-5}$ | 0.31<br>$p < 1 \times 10^{-5}$ | - |
| Other | 0.61<br>$p < 1.30 \times 10^{-4}$ | 0.34<br>$p < 1.80 \times 10^{-3}$ | 0.26<br>$p = 4.87 \times 10^{-3}$ | 0.47<br>$p < 1 \times 10^{-5}$ |

Average of the pairwise ERC scores between the proteins in different glycosylation pathways. Significance tested against 100,000 permutations.

### Not all glycosylation complexes have high correlated evolutionary rates

We next examined whether glycosylation proteins that form physical complexes have high ERC values. The GlcNAc transferase complex (GlcNAc-T) performs the first step in GPI anchor synthesis, attaching a GlcNAc to the phosphatidylinositol base (13). GlcNAc-T consists of seven proteins: PIGA, PIGC, PIGH, PIGP, PIGQ, PIGY, and DPM2. The proteins in the GlcNAc-T complex do not have enriched rate covariation with each other (mean ERC = 0.18, p = 0.41) (Figure 2A, S7 Fig A). Only one pair of proteins shows significant ERC, PIGQ and DPM2 (ERC = 3.58). The low ERC scores between proteins in this complex suggest that physical interaction does not drive evolutionary rate changes in these proteins. The other complex in GPI anchor synthesis is the GPI transamidase complex which is comprised of five proteins: PIGS, PIGT, PIGU, PIGK, and GPAA1 (13). The GPI transamidase complex attaches the protein to the synthesized GPI anchor. The mean ERC score among proteins in the GPI transamidase complex is 5.86 (p < 1×10^-5^), which is more than 11 standard deviations higher than the average expected based on permutation testing (permutations mean = 0.12) (Figure 2B, S7 Fig B). All pairwise ERC scores between GPI transamidase proteins show significant rate correlation (ERC ≥ 3), except between PIGU and PIGK (ERC = 0.80).

**Figure 2:**
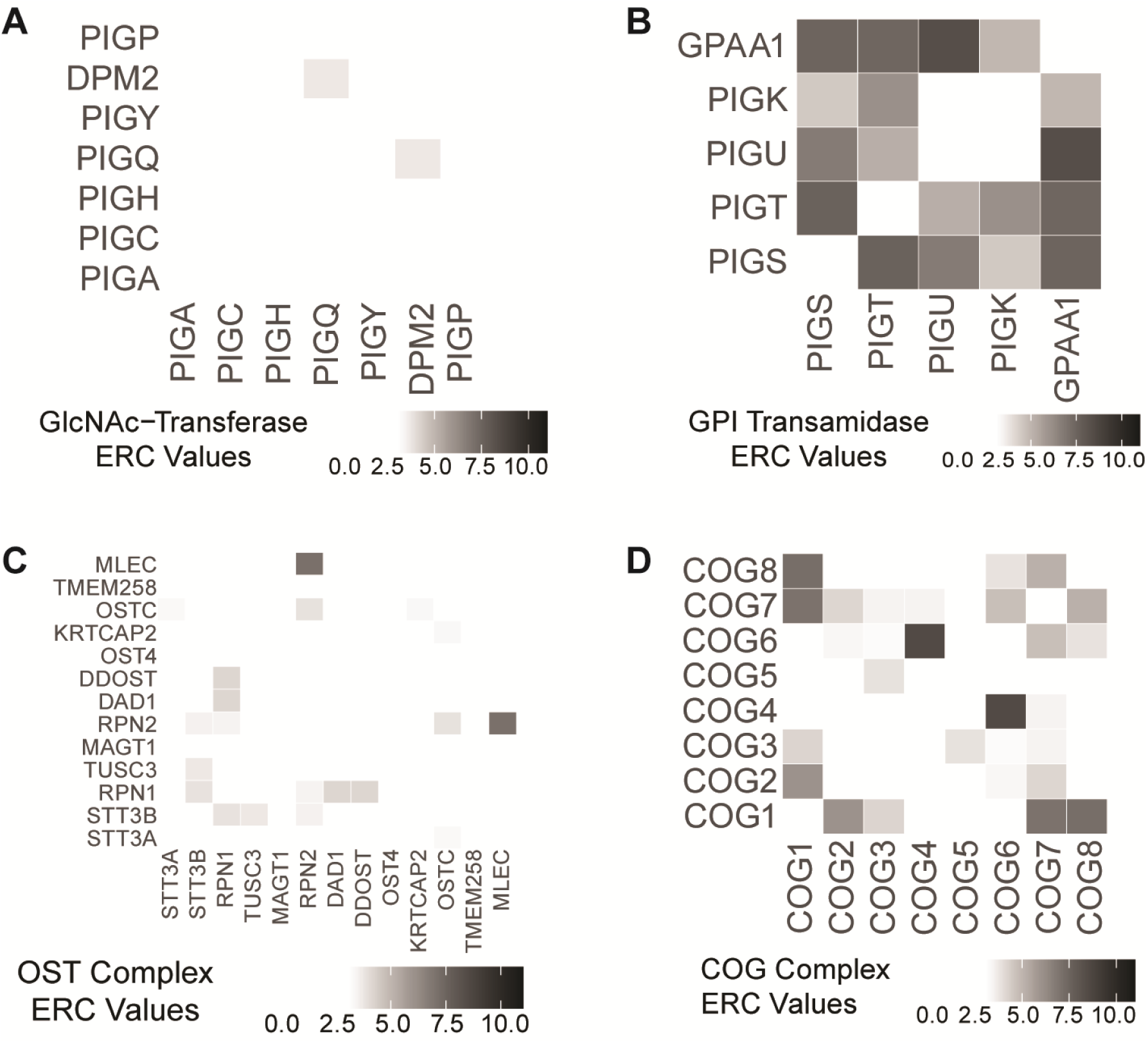
Heatmaps of significant pairwise ERC values for proteins in complexes involved glycosylation. The GlcNAc-T complex (A) does not show significant ERC (p = 0.41). The GPI transamidase complex (B), the OST complex (C), and the COG complex (D) all have significant ERC (p < 1×10^-^⁴).

In N-linked glycosylation, the oligosaccharyltransferase (OST) complex attaches the synthesized glycan onto a protein (11). This complex is made up of 13 proteins. The average pairwise ERC score of the OST complex is 1.26 and is higher than expected based on permutation testing (p = 4.0×10^-5^) (Figure 2C, S7 Fig C). The highest value is between MLEC and RPN2 (ERC = 7.02). Both MAGT1 and OST4 have no significant ERC scores with any other proteins in the complex. MAGT1 acts as a scaffold protein for STT3B in the OST complex, but it also functions as a magnesium transporter (21). Because MAGT1 has functions outside of this complex, it might have multiple evolutionary pressures that reduce its covariation rate with OST complex proteins. OST4 plays a critical role in the OST complex, binding the catalytic subunits STT3A or STT3B to the rest of the complex (22). Despite this, it does not show strong ERC with OST complex components suggesting that it might have another unknown, unrelated function.

The COG complex is a group of eight proteins (COG1-8) that acts as a tethering factor in intra-Golgi trafficking (23). Pairwise ERC scores between the proteins in the COG complex show significant ERC (mean ERC = 3.00, p < 1×10^-5^) (Figure 2D, S7 Fig D). Only half of the protein pairs show a significant correlation (ERC ≥ 3). The COG complex is organized into two lobes, with COG1-4 making up one lobe and COG5-8 making up the second lobe. Within the first lobe, COG1 has significant ERC with both COG2 and COG3. Strikingly, COG4 does not have any significant ERC scores with the other proteins in the same lobe; however, COG4 has the highest ERC value of the whole complex with COG6 (ERC = 8.46), a protein in the opposite lobe. The second lobe has slightly more protein pairs with significant ERC. COG6, COG7, and COG8 all have significant ERC with each other. COG5 has no significant ERC with other proteins in the same lobe. A bond between COG1 and COG8 connects the two lobes. COG1 and COG8 have the second highest ERC score of the complex (ERC = 7.16). The average ERC score between proteins in the first lobe is 2.70, and the average ERC score between proteins in the second lobe is 2.51. Unexpectedly, the average ERC score among inter-lobe protein pairs is 3.29. Strong ERC scores in this complex are not solely driven by direct physical contact between the proteins and are, on average, higher between proteins in different lobes.

### Glycosylation enzymes with similar functions have increased ERC

Next, we examined whether glycosylation proteins with similar enzymatic functions, but from different pathways display high ERC. There are twelve mannosyltransferases across the three glycosylation pathways. These mannosyltransferases are not known to physically interact with each other, and each one functions in only one specific glycosylation pathway. Pairwise ERC scores between these twelve mannosyltransferases show significant ERC by permutation testing (mean ERC = 3.07, p < 1×10^-5^) (Figure 3, S7 Fig E). Of these 55 ERC values, 27 are above the significance cut-off (≥3). The top score is between ALG12 and ALG2 (ERC = 8.02), which are both in the N-linked glycosylation pathway. The second highest score is between POMT1, involved in O-mannosylation, and ALG2, involved in N-linked glycosylation (ERC = 7.97). The six mannosyltransferases involved in N-linked glycosylation have an average ERC of 4.40. The four GPI anchor synthesis specific mannosyltransferases have an average of 2.32. O-linked glycosylation has only two mannosyltransferases (POMT1 and POMT2), and their average ERC score is 6.86.

**Figure 3:**
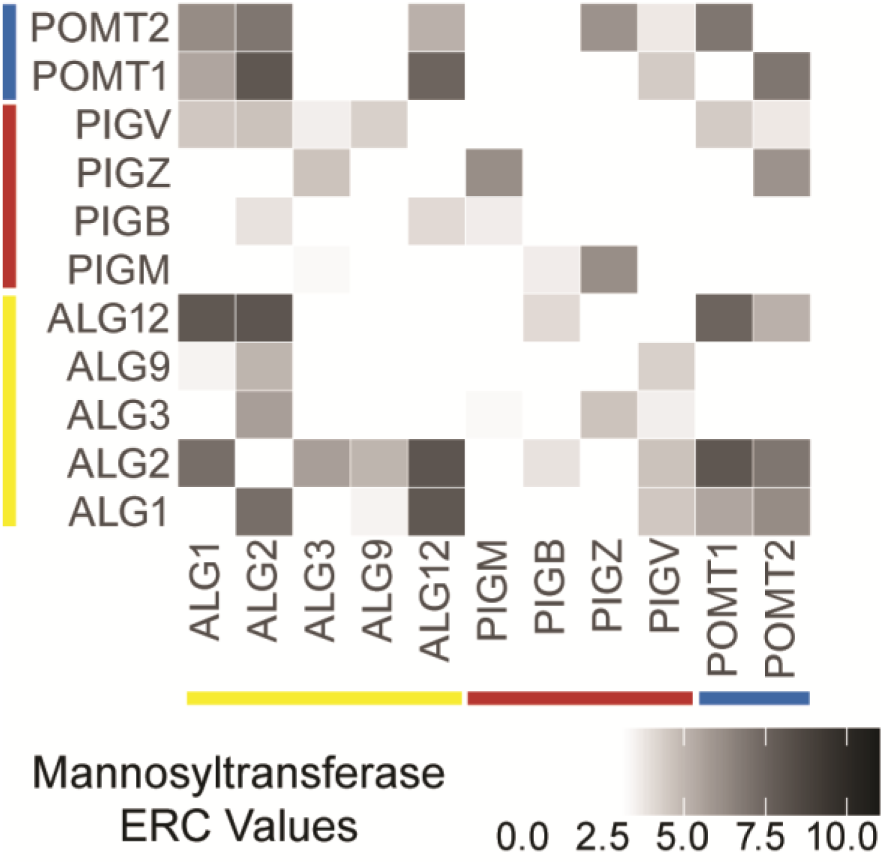
Heatmap showing significant ERC values between mannosyltransferases. N-linked glycosylation (yellow), GPI-anchor synthesis (red), O-linked glycosylation (blue). Mannosyltransferases have significant ERC with each other across pathways (p < 1×10^-^⁵).

### Glycosylation protein pairs with strongest ERC

Some of the top ten highest-scoring glycosylation protein pairs show overlapping functions, which could explain the high ERC scores (Table 3). For example, COG3 and TRIP11 (ERC = 9.62) are proteins bound to the outer membrane of the Golgi involved in vesicle tethering and assembly and maintenance of the Golgi. PIGZ and PIGW also have a high ERC score (ERC = 9.10). PIGZ and PIGW are both involved in GPI anchor synthesis in the ER, and they share a common GPI recognition domain in their transmembrane domains (24). While these two pairs have high ERC scores supported by current knowledge, the top scores also include pairs with high scores that are not easily explained based on our current knowledge of their function. FKRP, an enzyme that adds ribitol to O-linked alpha-dystroglycans in the Golgi (25), and MOGS, an enzyme that cleaves glucose from the synthesized N-glycan in the ER (26), have an ERC score of 10.63. EXTL3, a GlcNAc transferase for O-linked glycosaminoglycan synthesis located in the Golgi (27), and GMPPA, a proposed inhibitor of GDP-mannose synthesis localized to the cytoplasm (28), have an ERC score of 9.62. The remaining six pairs in the top 10 show similar patterns of high ERC with no obvious overlap in function, localization, or glycosylation pathway. These unrelated pairs of proteins with high ERC values could signal novel interactions between glycosylation proteins.

**Table 3:**
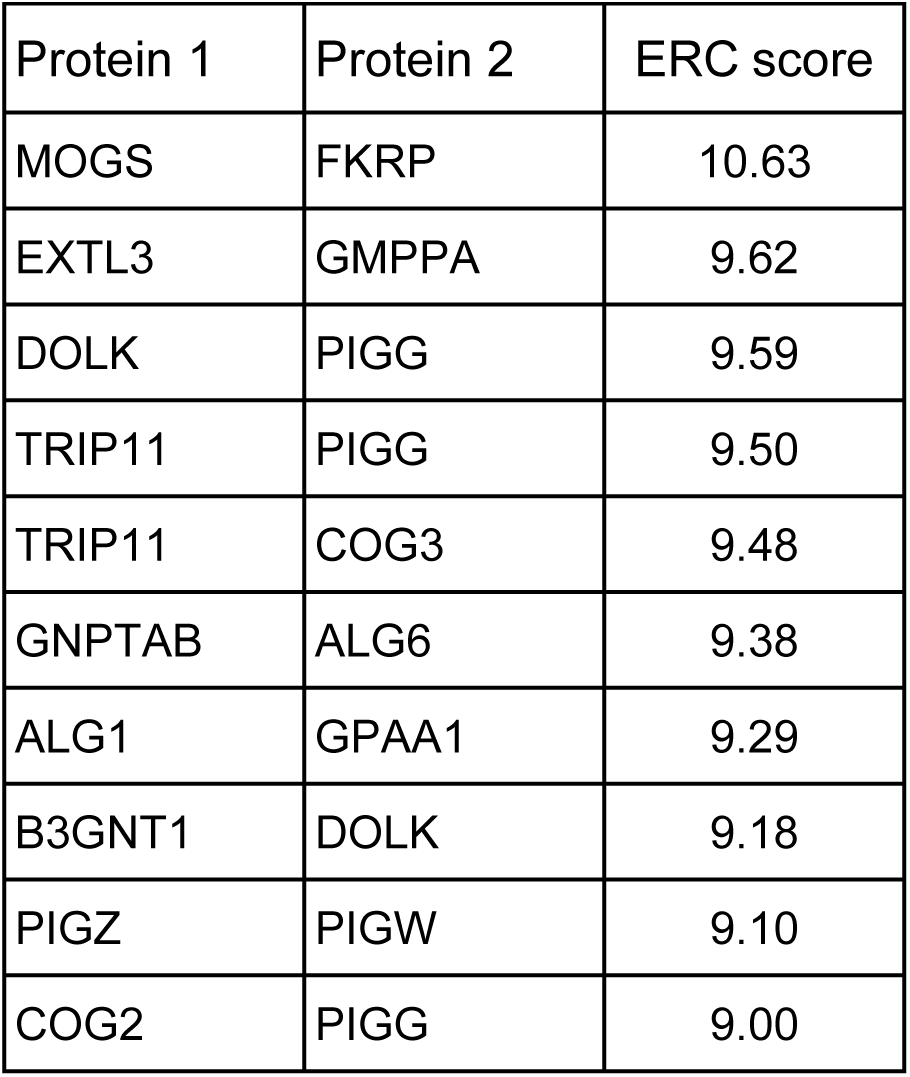
Top 10 pairwise ERC scores among glycosylation proteins.

### Genome-wide patterns of evolutionary rate correlation for GPI Anchor Synthesis Proteins

We next examined whether proteins with unrelated, non-glycosylation functions might also have high ERC values with glycosylation proteins. To do this, we extended our analyses to a full genome-wide comparison. Because the GPI anchor synthesis pathway had the highest average ERC score of all glycosylation pathways (GPI mean ERC = 1.61), we focused the genome-wide analyses on the GPI anchor biosynthesis pathway. We included the 27 proteins specific to the GPI anchor synthesis pathway and DPM1, DPM2, and DPM3. DPM2 is a component of the GlcNAc-transferase complex in GPI anchor synthesis and the dolichol-phosphate-mannose (DPM) synthase complex with DPM1 and DPM3. DPM synthase is required to add mannose to the GPI anchor (29). We calculated pairwise ERC values for each of the 30 proteins involved in GPI anchor biosynthesis with all 19,149 proteins in our dataset (Figure 4, S1 Data). This resulted in 568,438 pairwise values ranging from -8.97 to 11.28, with an average of 0.32. 31,483 protein pairs had significant ERC scores (ERC ≥ 3).

**Figure 4:**
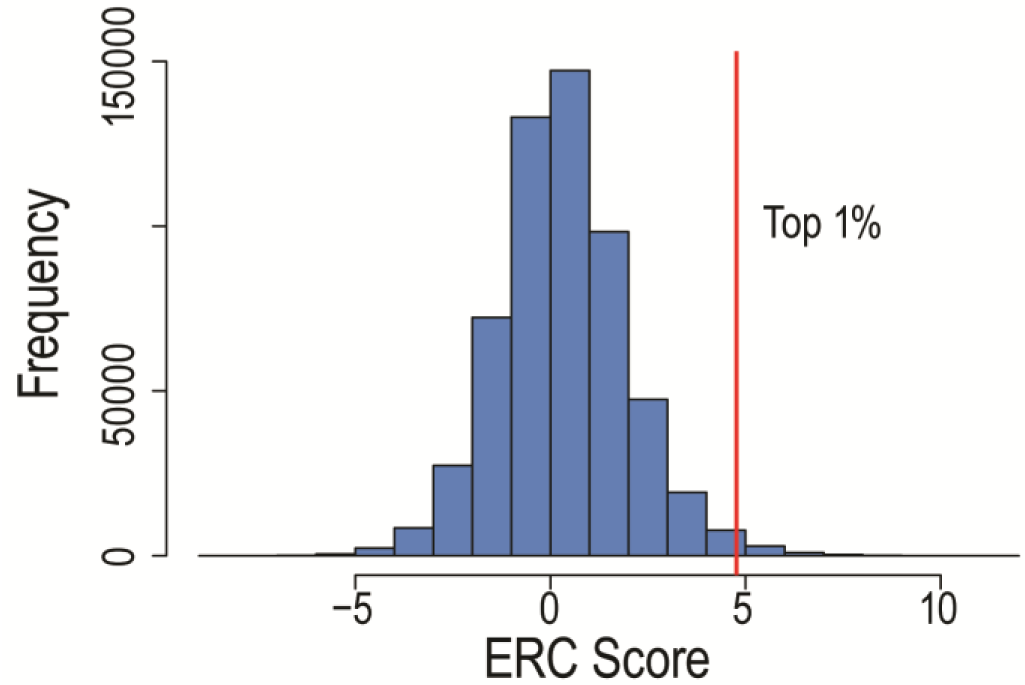
Histogram of genome-wide ERC values for with GPI anchor synthesis proteins. Vertical red line indicates top 1% of values.

### Gene Ontology enrichment of proteins showing high ERC with GPI Anchor Synthesis

Gene Ontology (GO) analysis of the top 1% of GPI anchor synthesis genome-wide ERC values (5685 values, ERC > 4.77, 3386 proteins) (Figure 5, S2 Data) revealed enrichment in two glycosylation related pathways, *GPI anchor biosynthetic process GO:0006506* (fold enrichment = 3.86, q = 1.03×10^-3^) and *dolichol-linked oligosaccharide biosynthetic process GO:0006488* (fold enrichment = 4.70, q= 1.36×10^-3^), which support the finding described above that GPI anchor synthesis proteins show strong signals with themselves and other glycosylation pathways.

**Figure 5:**
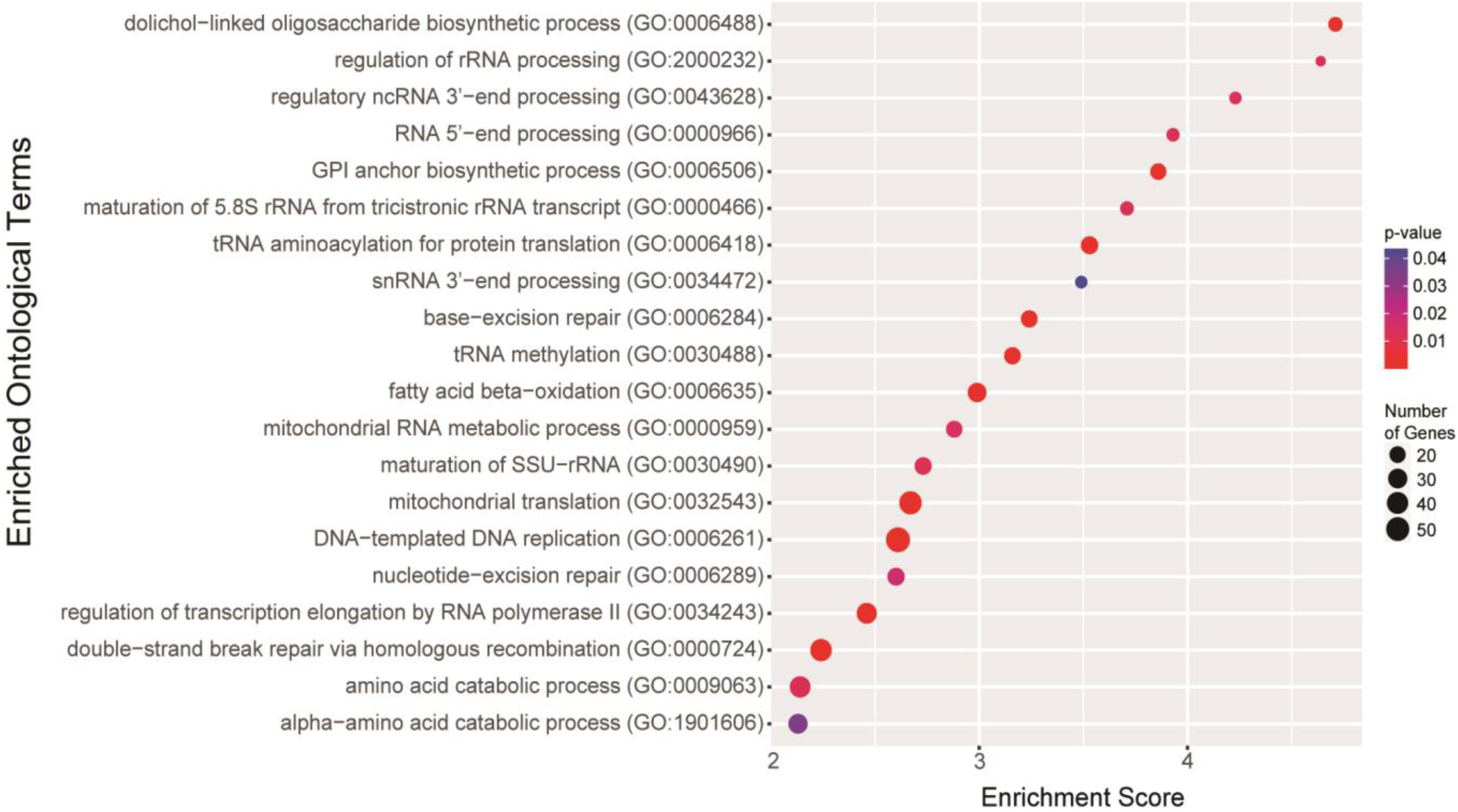
GO terms from analysis of proteins in the top 1% of GPI anchor synthesis genome-wide ERC values. P-values are indicated by red-to-blue gradient, with red the lowest p-values and blue the highest p-values. Gene number identified in each category is indicated by the size of the circle.

GO analysis also returned multiple terms relating to ncRNA modification: *regulation of rRNA processing GO:2000232* (fold enrichment = 4.63, q = 1.37×10^-2^), *regulatory ncRNA 3’-end processing GO:0043628* (fold enrichment = 4.23, q = 1.34×10^-2^), and *tRNA methylation GO:0030488* (fold enrichment = 3.16, q = 3.48×10^-3^) (Figure 5, S2 Data). Previous work from our lab also identified ncRNA processing proteins as potential modifiers of NGLY1-CDG (15). The enrichment of ncRNA processing proteins in this analysis further supports an unexplored connection between glycosylation proteins and ncRNAs. There were also multiple GO terms related to DNA repair: *base excision repair GO:006284* (fold enrichment = 3.24, q = 2.71×10^-3^), *nucleotide-excision repair GO:0006289* (fold enrichment = 2.60, q = 1.83×10^-2^), and *double-strand break repair via homologous recombination GO:0000724* (fold enrichment = 2.24, q = 2.27×10^-3^). This enrichment was unexpected as no previous evidence links DNA repair to GPI anchor synthesis and suggests a possible interaction between the two pathways.

Loss of function in most GPI anchor synthesis proteins leads to a CDG. To identify potential proteins that could modify multiple CDG genes across the pathway, we also ran GO analysis on proteins that had a significant ERC score with at least a third (10 or more) of the 30 GPI anchor synthesis proteins (Figure 6, S2 Data). Unsurprisingly, GO analysis again showed high enrichment for glycosylation proteins (*GPI anchor biosynthetic process* GO:0006506, fold enrichment 18.53, q = 1.59×10^-4^; *dolichol-linked oligosaccharide biosynthetic process* GO:0006488, fold enrichment 15.63, q = 4.48×10^-2^) as well as terms related to ncRNA (*tRNA aminoacylation for protein translation GO:0006418*, fold enrichment = 15.63, q = 8.10×10^-5^; *tRNA methylation GO:0030488*, fold enrichment = 11.45, q = 7.69×10^-3^; *rRNA processing GO:0006364*, fold enrichment = 4.52, q = 7.26×10^-3^). Unlike what was found when analyzing the top 1% of values, there was also enrichment for ribosome assembly (*ribosomal small subunit biogenesis GO:0042274*, fold enrichment = 8.51, q = 9.95×10^-5^; *ribosome assembly GO:0042255,* fold enrichment = 7.46, q = 4.73×10^-2^) and m*itotic recombination GO:0006312* (fold enrichment = 17.27, q = 3.43×10^-2^). These pathways interacting with multiple proteins in GPI anchor synthesis may represent compelling therapeutic targets for multiple GPI anchor deficiencies.

**Figure 6:**
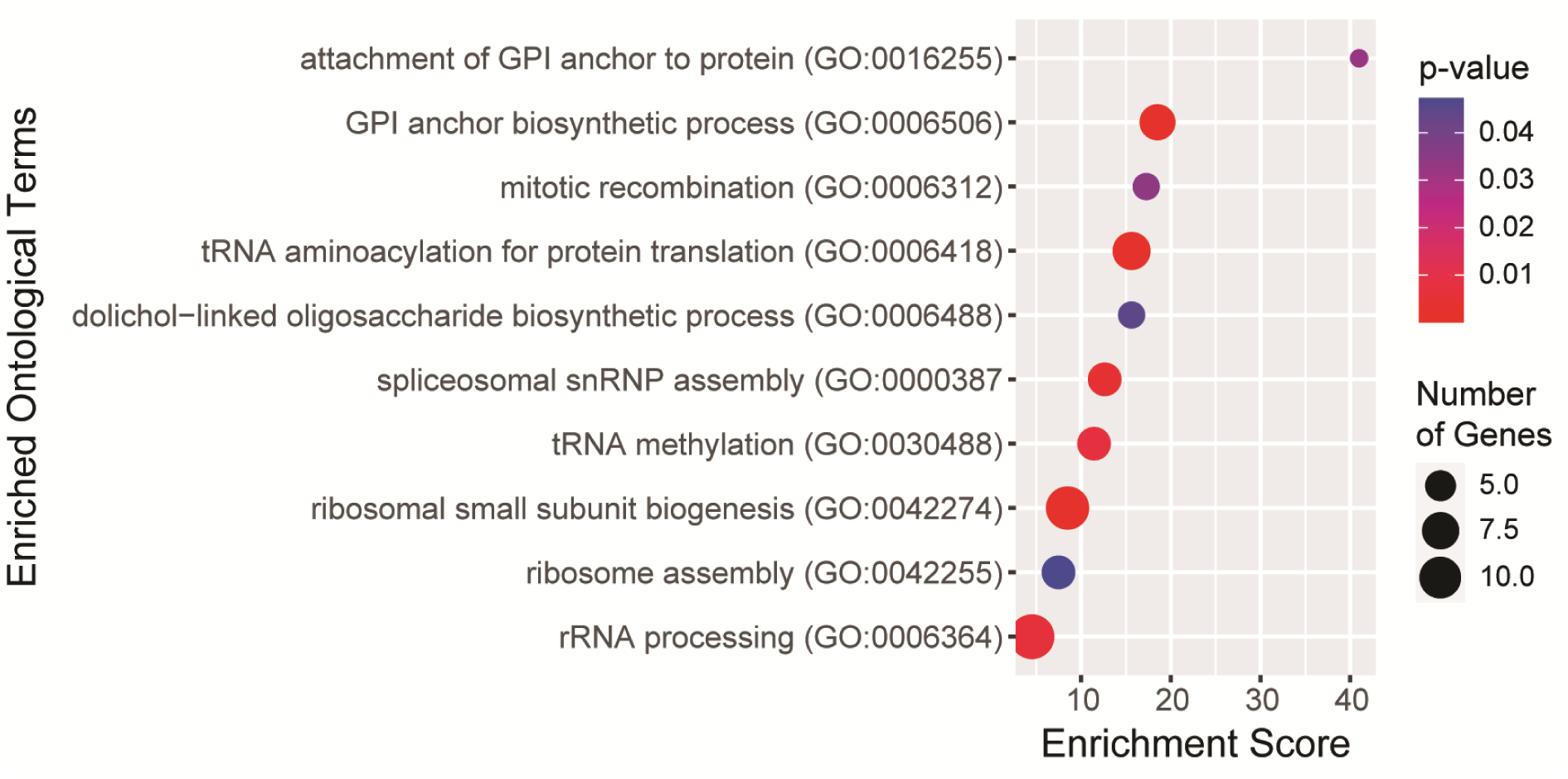
GO terms from analysis of proteins with significant ERC values with at least 10 GPI anchor synthesis proteins. P-values are indicated by red-to-blue gradient, with red the lowest p-values and blue the highest p-values. Gene number identified in each category is indicated by the size of the circle.

### Top scoring pairs across the genome

Many top-scoring pairs from the genome-wide analysis fall under the categories enriched in the GO analyses. Four of the top 30 scoring pairs were between known glycosylation proteins (Table 4). PIGN, a phosphoethanolamine transferase (13), had high ERC with MAN2B2, a protein involved in glycan recycling (ERC = 10.01) (10). PIGG, a phosphoethanolamine transferase (13), had strong ERC with DOLK, a kinase that creates dolichol-phosphate upon which the N-glycan is built (ERC = 9.59) (30), and TRIP11, a protein involved in Golgi vesicle tethering (ERC = 9.50) (31). GPAA1, a member of the GPI transamidase complex (13), had high ERC with ALG1, a mannosyltransferase in N-linked glycosylation (ERC = 9.29) (11). All seven glycosylation proteins in these top-scoring pairs are associated with their own CDGs, further suggesting that known CDG genes are likely candidates for genetic modifiers of other CDGs (14–16,20).

**Table 4:** Top scoring protein pairs from genome-wide analysis of GPI anchor synthesis proteins.

| Rank | GPI Protein | Protein 2 | ERC score |
| --- | --- | --- | --- |
| 1 | PIGG | HTT | 11.28 |
| 2 | PIGQ | WDR90 | 10.50 |
| 3 | PIGG | TUBGCP2 | 10.49 |
| 4 | PIGG | ATG7 | 10.44 |
| 5 | PIGB | UTP20 | 10.33 |
| 6 | PIGW | DHODH | 10.32 |
| 7 | PIGG | TTC8 | 10.22 |
| 8 | PIGO | ZNF622 | 10.19 |
| 9 | PIGB | AP4E1 | 10.09 |
| 10 | PIGG | PRKDC | 10.01 |
| 11 | PIGN | MAN2B2 | 10.01 |
| 12 | GPAA1 | MCM7 | 9.97 |
| 13 | PIGG | ERCC6 | 9.76 |
| 14 | PIGN | WDR36 | 9.74 |
| 15 | PIGG | DOLK | 9.59 |
| 16 | PIGA | ZFYVE20 | 9.58 |
| 17 | PIGZ | C12orf65 | 9.57 |
| 18 | PIGB | GEMIN5 | 9.57 |
| 19 | PIGG | TRIP11 | 9.50 |
| 20 | PIGG | GTF3C5 | 9.47 |
| 21 | PIGM | INF2 | 9.45 |
| 22 | PIGQ | PKD1 | 9.40 |
| 23 | PIGN | CTDP1 | 9.39 |
| 24 | PGAP4 | DHX40 | 9.38 |
| 25 | PIGG | PHRF1 | 9.36 |
| 26 | PIGB | MSH6 | 9.33 |
| 27 | PIGO | SHPRH | 9.31 |
| 28 | GPAA1 | ALG1 | 9.29 |
| 29 | PIGG | AP5Z1 | 9.26 |
| 30 | PGAP1 | SMYD3 | 9.22 |

Top scores in our dataset revealed high ERC between PIGB and UTP20 (ERC = 10.33) and PIGN and WDR36 (ERC = 9.74). UTP20 and WDR36 are both involved in rRNA processing (32,33). Top scores also included PIGG and PRKDC (ERC = 10.01), a DNA damage sensing protein (34), ERCC6 (ERC = 9.76), a protein important for transcription-coupled excision repair (35), and AP5Z1 (ERC = 9.26), a protein involved in homologous recombination DNA repair (36). GPAA1 had high ERC with MCM7 (ERC = 9.97), a protein that signals DNA damage (37). These top pairs further support the hypothesis of a relationship between GPI anchor synthesis genes and the GO terms identified and offer targeted protein pairs to begin testing to explore these pathways.

### Cytoplasmic GPI anchor synthesis proteins have lower ERC

Among the top 1% of genome-wide GPI anchor scores, nearly 10% are with PIGG (552/5685). To determine if this enrichment was significant, we compared the actual number of ERC values each GPI anchor synthesis protein had in the top 1% to an expected number of values, assuming an equal distribution of all the GPI anchor synthesis proteins (expected ∼189 occurrences per protein) (Figure 7A, S1 Table). PIGG (552 occurrences, 2.91 fold change, p < 2.2×10^-16^) was the most overrepresented, with nearly three times as many ERC values in the top 1% as expected, followed by PIGW (526 occurrences, 2.78 fold change, p < 2.2×10^-16^). On the other hand, PIGY (8 occurrences, -23.68 fold change, p < 2.2×10^-16^), PIGH (7 occurrences, -27.06 fold change, p < 2.2×10^-16^), and DPM2 (6 occurrences, -31.57 fold change, p < 2.2×10^-16^) all have more than 20 fold less than the expected frequency. We tested whether this enrichment/depletion followed the pathway order (Figure 7B, 7C). Analyses of the GPI anchor synthesis proteins by pathway order demonstrated that 8/10 proteins on the cytoplasmic side are underrepresented, and none are overrepresented. In the ER lumen, only 4/17 proteins are underrepresented, and 9/17 of the proteins are overrepresented.

**Figure 7:**
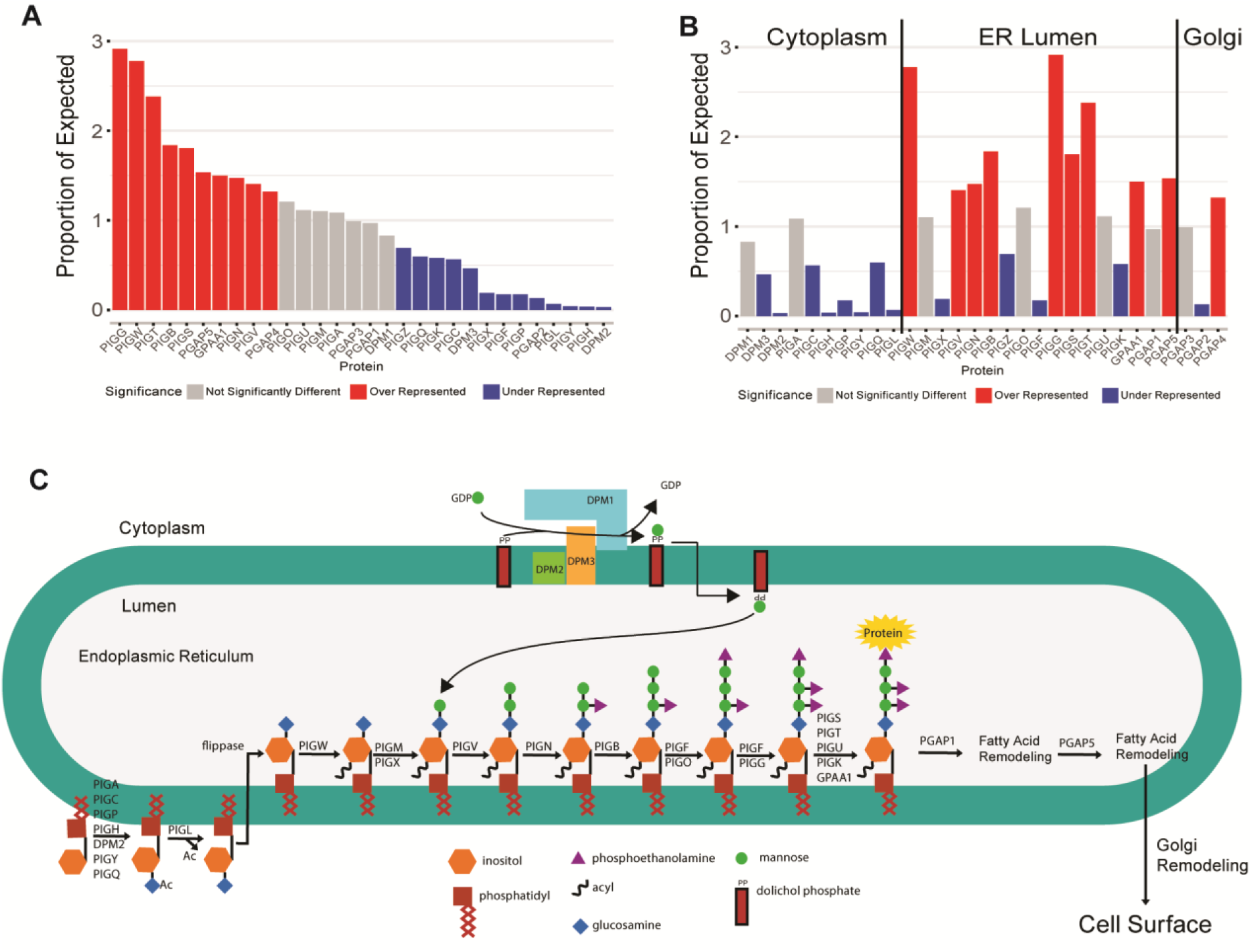
Proportion of expected occurrences of all GPI anchor synthesis proteins in the top 1% of genome-wide scores. Overrepresented proteins are shown in red, underrepresented in blue, and proteins not significantly different from expected are in grey. A) Proteins sorted from most to least represented. B) Proteins sorted in pathway order with lines dividing genes active on the cytoplasmic side of the ER, the lumen side of the ER, and in the Golgi. C) GPI anchor synthesis pathway diagram. Synthesis begins on the cytoplasmic side of the ER. The anchor is flipped to the lumen side. After the protein is attached, the glycan is sent to the Golgi for lipid remodeling.

Proteins acting on the cytoplasmic side of the ER were significantly underrepresented in the top 1% of ERC values, and proteins acting on the lumen side were highly overrepresented. We hypothesized that high ERC with other ER resident proteins may be driving the overrepresentation of lumen proteins. However, GO analysis for cellular compartment of the top 1% of proteins with high ERC with GPI anchor synthesis proteins revealed the top enriched compartments are associated with the exosome, ribosome, and mitochondria (S8 Fig, S2 Data). Some ER-related terms were enriched, but over 50 other terms were more enriched. This indicates that while ER resident proteins may contribute to some of the high scores observed among lumen GPI anchor synthesis proteins, it is unlikely that high scores with ER resident proteins are solely responsible for driving the overrepresentation of lumen GPI anchor synthesis proteins in the top 1% of ERC scores.

We examined the N-linked glycosylation pathway to determine if lower ERC scores for cytoplasmic enzymes were unique to GPI anchor synthesis or common in glycosylation pathways. The N-linked glycosylation pathway is similar in that the first few steps occur on the cytoplasmic side, and the glycan is completed and attached to a protein on the ER lumen side. We calculated ERC scores for proteins in the N-linked glycosylation pathway across the genome and ran the same analysis as we did for GPI anchor synthesis. We took the top 1% of values and determined the number of scores for each N-linked glycosylation protein and compared this to the expected distribution, as we did above (S9 Fig). Among the eight cytoplasmic proteins, three were under-represented, and four were over-represented. Among the ER lumen proteins, eight of the 23 proteins were under-represented, and ten of the 23 lumen proteins were over-represented. The N-linked glycosylation enzymes acting on the cytoplasmic did not have an enrichment of lower scores as observed for GPI anchor synthesis. The ERC pattern for cytoplasmic vs. ER lumen proteins appears specific to the GPI anchor synthesis pathway.

### *In vivo* testing of top ERC pairs confirms genetic interactions

A strong ERC score is an indicator of shared function between proteins in a complex or across a pathway (19). Therefore, we tested top ERC scoring protein pairs to find new genetic interactions with GPI anchor synthesis proteins (Table 5, S3 Data). Because many GPI anchor synthesis proteins are necessary for survival in *Drosophila*, ubiquitous loss-of-function experiments are challenging. The *Drosophila* eye provides a powerful alternative model for testing genetic interactions with glycosylation genes (14). Using the GAL4-UAS system with the *eyes absent* composite GAL4 driver (*eya*) (38), we expressed RNAi in an eye-specific manner to create knockdown models for each GPI anchor gene tested. Many of these GPI anchor knockdown models have changes in eye size/quality/color. We can observe genetic interactions by creating double knockdowns of the highest scoring protein pairs in our genome-wide list and comparing the effects of each gene’s single knockdown with the effects of the double knockdown.

**Table 5:** *In vivo* testing for genetic interactions of top scoring protein pairs.

|  |  |  |  |  |  |
| --- | --- | --- | --- | --- | --- |
|  | GPI protein | Protein 2 | Protein 2 function | ERC score | Interaction |
| GPI x CDG | PIGG | DOLK | Dolichol-phosphate synthesis involved in N-linked glycosylation | 9.59 | YES |
|  | PIGG | TRIP11 | Golgi maintenance and homeostasis | 9.50 | NO |
|  | GPAA1 | ALG1 | First mannosyltransferases in N-linked glycosylation | 9.29 | YES |
| GPI x Genome-wide | PIGA | RBSN | Early endocytic membrane fusion and membrane trafficking of recycling endosomes | 9.58 | YES |
|  | PIGW | DHODH | Pyrimidine synthesis | 10.32 | NO |
|  | PIGM | INF2 | Actin polymerization and depolymerization | 9.45 | YES |
|  | PIGZ | C12orf65 | Mitochondrial gene translation | 9.57 | NO |
|  | PIGO | ZNF622 | Ribosomal maturation | 10.19 | YES |
|  | PIGO | SMPD4 | Sphingomyelin catabolism to form phosphorylcholine and ceramide | 9.13 | YES |
|  | PIGG | HTT | Vesicle transport | 11.28 | NO |
|  | PIGG | TUBGCP2 | Microtubule nucleation | 10.49 | YES |
|  | PIGG | ATG7 | Autophagy | 10.44 | NO |
|  | PIGG | TTC8 | Cilia membrane protein sorting | 10.22 | YES |
|  | PIGG | PHRF1 | Protein ubiquitination | 9.36 | NO |
|  | PIGG | ZGPAT | Transcription repressor, negatively regulates EGFR expression | 9.18 | YES |
|  | PIGT | HDHD3 | Function unknown | 9.20 | YES |

Top scoring protein pairs tested for interaction in a *Drosophila* eye model. GPI proteins sorted by pathway order. Interactions tested with two-way ANOVA $p < 0.05$ .

This tissue-specific eye model provides an efficient way to screen for genetic interactions. However, there are caveats in using an eye knockdown model. Most genetic interactions are context-specific. Because we are knocking down the genes only in the eye, we may not be testing in the correct tissues for the interactions identified by ERC. All the genes tested have expression in the *Drosophila* eye (39), but the eye does not always have the highest tissue expression. Another major caveat to this model is the use of RNAi to knockdown the genes of interest. Different RNAi transgenes reduce the expression level of their target genes by different levels (S3 Data). It is possible that the interactions in our models require specific expression levels that a particular RNAi does not achieve.

We tested the top sixteen pairs from our genome-wide list with available RNAi lines (Table 5, S10 Fig, S3 Data). Of the sixteen pairs tested, 10 showed significant interactions. With over half of the tested pairs validating, this is a higher hit rate than typically seen with ERC (40–42). When multiple RNAi lines were available for a gene, we tested all RNAi lines. qPCR was performed on all RNAi lines to determine knockdown levels for each line. The RNAi line resulting in the lowest expression showed the interaction for almost all pairs with multiple RNAi lines tested. In two interaction pairs, the lower knockdown RNAi line showed interaction while the strongest knockdown line did not. This indicates expression level may play a role in identifying genetic interactions. Here we highlight two interactions: one pair between a GPI anchor synthesis protein and another glycosylation protein, and one pair between a GPI anchor synthesis protein and a non-glycosylation protein.

GPAA1 and ALG1 had one of the highest scores between a GPI anchor synthesis protein and a glycosylation protein outside of GPI anchor synthesis (ERC = 9.29). GPAA1 is a catalytic component of the GPI-transamidase complex that attaches a protein to the synthesized GPI anchor (13,43). ALG1, located on the cytoplasmic side of the ER, adds the first mannose onto the N-linked glycan (11). Both *GPAA1* and *ALG1* are associated with their own respective CDGs (44). While eye-specific knockdown of *ALG1* alone resulted in no change in eye size, it did result in structural disorganization and occasional necrosis. Eye-specific knockdown of *GPAA1* alone resulted in a significant reduction in eye size, but there was little qualitative change (Figure 8 A, B). Double knockdown of *GPAA1* and *ALG1* resulted in a significantly smaller eye than the expected additive effect of both single knockdowns (p = 9.91×10^-5^), and the eye was structurally disorganized. This indicates a genetic interaction between *GPAA1* and *ALG1*, two genes involved in distinct glycosylation pathways.

**Figure 8:**
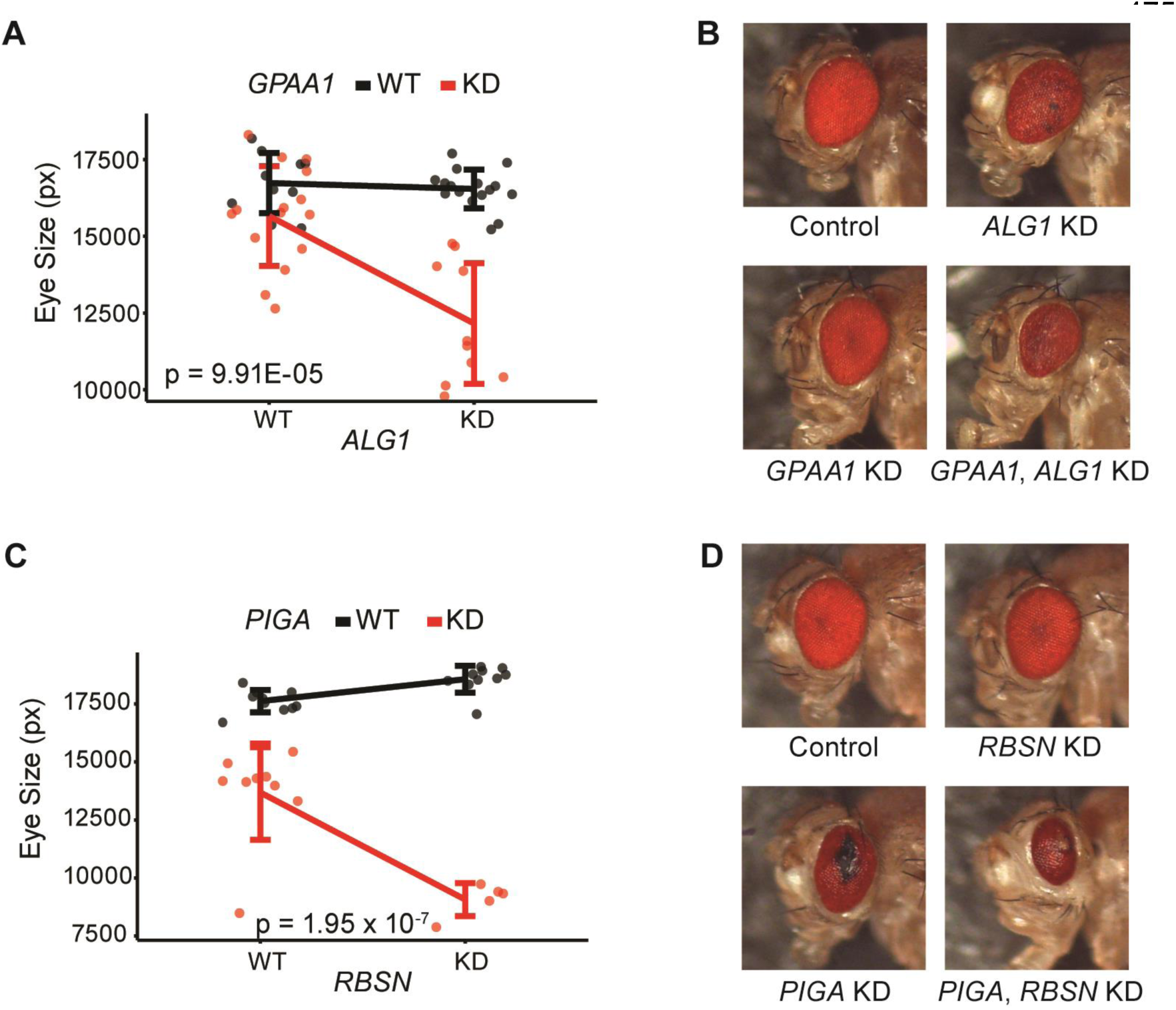
*In vivo* testing for genetic interactions in the eye. A),C) Quantification of eye sizes. Quantification in pixels (px) is along the Y axis. X axis has expression level of Protein 2, Wild Type (WT) or Knockdown (KD), and line color indicates expression level of GPI anchor synthesis protein (WT or KD). Interaction p-value calculated using a two-way ANOVA. B),D) Representative images of *Drosophila* eyes showing WT control, single KDs, and double KD eyes.

RBSN has the top ERC score with PIGA (ERC = 9.58), the enzymatic component of GlcNAc-transferase in GPI anchor synthesis (13). RBSN is recruited to the early endosome and plays a role in sorting cargo between lysosomal degradation and membrane recycling (45,46). While single knockdown of *RBSN* in the eye yields no change in eye size or structure (Figure 8 C, D), single knockdown of *PIGA* results in a significantly decreased eye size. *PIGA* knockdown also has structural disorganization, necrosis, and glassiness in the eye. Double knockdown of *PIGA* and *RBSN* enhances the small eye *PIGA* phenotype in a non-additive manner indicating a genetic interaction between *PIGA* and *RBSN* (p = 1.95 x 10^-7^). The double knockdown continues to show necrotic tissue and disorganization. This same interaction was seen with a second RNAi line for *PIGA*, where the loss of *RBSN* enhances the phenotype, in a non-additive manner (p = 2.70 x 10^-4^) (S10 Fig D).

### High ERC scores suggest a connection between GPI anchor synthesis and the endosome

It may not be immediately apparent why GPI anchor synthesis proteins like PIGA might show strong ERC and a genetic interaction with endosomal proteins like RBSN. However, many GPI-anchored proteins are frequently endocytosed and are either recycled back to the cell membrane or sent to the lysosome for degradation (47). RBSN is recruited to the early endosome by Rab5 where it aids in vesicle fusion to form the early endosome (46). Interestingly, Rab5 also binds HTT (48). HTT is an essential protein implicated in many cell functions, including cell transport and autophagy, and had the highest genome-wide ERC score with a GPI anchor synthesis protein (HTT and PIGG, ERC = 11.28) (49). Many interactors have been identified for HTT, but none are involved in GPI anchor synthesis. Because of the genetic interaction between RBSN and PIGA and the link between RBSN, Rab5, and HTT, we further examined the evolutionary covariation between RBSN, HTT, and the GPI anchor pathway.

RBSN and HTT show significant ERC scores with 15 and six of the 30 GPI anchor synthesis proteins, respectively. Strikingly, all six GPI anchor synthesis proteins, including PIGG, that had high ERC scores with HTT also had high scores with RBSN (PIGG with RBSN, ERC = 9.05; PIGG with HTT, ERC = 11.28) (green proteins in Figure 9). HTT and RBSN also had a very high ERC score with each other (ERC = 8.21). These high ERC scores suggest a previously unknown functional connection between GPI anchor synthesis proteins, HTT, and RBSN, and more broadly, with the endosome pathway. This also highlights the need to examine all proteins in the GPI anchor synthesis pathway together. A high ERC score between HTT and PIGG or between PIGA and RBSN is individually interesting but challenging to contextualize. However, the fact that the majority of GPI anchor synthesis pathway proteins show high ERC with RBSN, which is directly linked to HTT, allows for better contextualization of the top hit in the genome-wide data.

**Figure 9:**
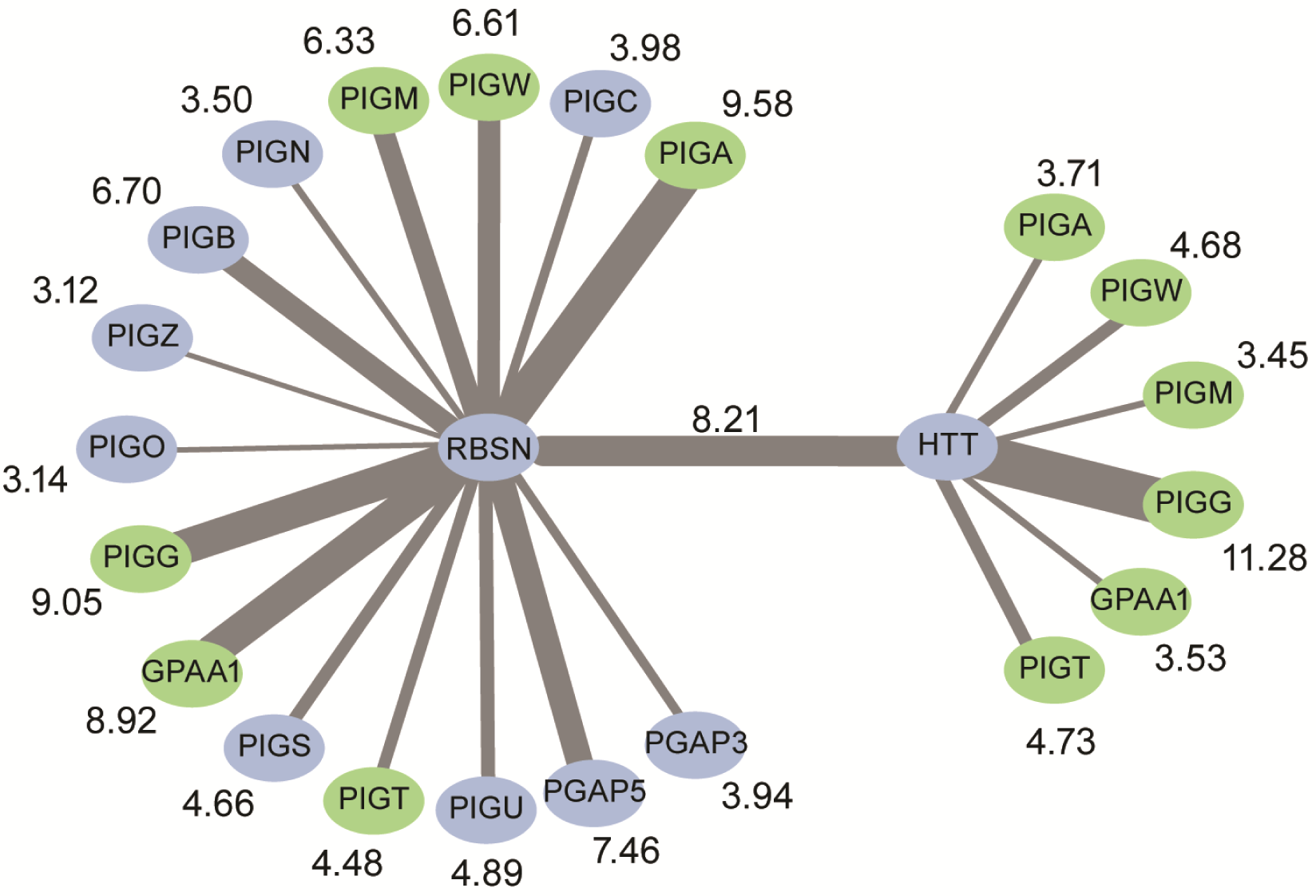
RBSN and HTT show strong ERC with each other and proteins in the GPI anchor synthesis pathway. ERC scores between proteins are indicated by the width of the edges and shown next to the GPI anchor protein. GPI anchor synthesis proteins showing significant ERC with both HTT and RBSN are shown in green. GPI anchor synthesis proteins are listed in pathway order.

## DISCUSSION

CDGs are poorly understood, multi-systemic disorders with extensive phenotypic variability, suggesting a role for genetic background and modifier genes. Identifying genetic modifiers of CDGs could help clarify the pathology of these disorders. In this study, we used evolutionary methods to explore genetic modifiers of CDGs and prioritize candidate genes for *in vivo* testing.

Until now, ERC has been used primarily to characterize unknown proteins or identify new proteins involved in specific pathways (19,40–42,50–53). One previous study used evolutionary similarities between disease genes to prioritize novel disease-causing genes (54). Because modifier genes genetically interact with their disease genes, it is logical that they would also be evolutionarily related to each other. Glycosylation is highly conserved from yeast to mammals allowing for strong evolutionary signals to have developed over time (11,55,56). Previously identified modifier genes for CDGs have high ERC values, including PMM2 – PGM1 (7.08) and DPAGT1 – DPM1 (4.17) (14,16). *In* vivo testing of top hits from our analysis had a hit rate (10 out of 16) higher than previously found in other ERC studies (40–42). Using ERC to identify disease modifiers is a novel application that could be applied to help better understand and treat rare diseases.

On average, glycosylation protein pairs showed high ERC regardless of whether the two proteins were in the same pathway. However, we found exceptionally high ERC scores between proteins involved in GPI anchor synthesis and N-linked glycosylation. One hypothesis for this might be both pathways’ shared reliance on mannose. N-glycans and GPI anchors have multiple mannoses added to their glycan chains, and they both rely on the DPM complex to supply mannose into the lumen of the ER (11,29). Only one type of O-linked glycan incorporates mannose, which could explain why O-linked glycosylation does not have as high of ERC scores with the other glycosylation pathways (57). Another potential explanation could be the necessity of N-linked glycosylation for proper folding of GPI-anchored proteins (58). In the ER, N-glycans on GPI-anchored proteins allow them to interact with chaperone proteins for proper protein folding. This requirement for both N-linked glycosylation and GPI anchor synthesis for protein folding might explain why the two different glycosylation pathways are evolutionarily linked. High ERC between N-linked glycosylation and GPI anchor synthesis could also suggest unexplored connections between these pathways, and they may need to be studied together instead of in isolation.

High ERC scores occur when two proteins are under similar evolutionary constraints (19,59). When analyzing ERC scores of GPI anchor synthesis proteins across the genome, we found that GPI anchor synthesis proteins acting on the cytoplasmic side of the ER had fewer high ERC scores than proteins acting on the luminal side of the ER. This was unexpected because all the proteins in the GPI anchor synthesis pathway are only known to act in that pathway. It is feasible that cytoplasmic components interact with a wider variety of proteins, which also influence their evolutionary rates in different ways, making it more difficult for them to have strong ERC scores. Cytoplasmic GPI anchor synthesis proteins may perform more functions than are currently known.

GO analysis of proteins showing high ERC with GPI anchor synthesis proteins resulted in multiple terms related to ncRNA. Until recently it was thought there was no overlap between RNA and glycosylation. In 2021, a group isolated glycosylated ncRNAs (60). It is currently unknown how these glycans are synthesized and attached to RNA (61). It is possible the same enzymes responsible for building the glycans for protein glycosylation also build RNA glycans. This relationship could explain the evolutionary connection between ncRNAs and glycosylation proteins.

Among the ncRNA GO terms, many were related to rRNA biology. However, it is unclear how glycosylation is involved in rRNA biology. Since glycosylation is a co-translational/post-translational modification, it suggests the evolutionary relationship identified between glycosylation proteins and rRNA processing proteins could be due to glycosylation’s dependence on the ribosome functioning properly and translating proteins (62). Further studies need to be performed to identify the specific protein interactions driving the association between these two pathways.

CDG patients have a broad clinical spectrum influenced by different disease-causing variants, environmental factors, and genetic background (63,64). Previous attempts to identify genetic modifiers with standard methods (i.e., mutagenesis screens and using natural variation in the population) have found promising candidates (14–16,20). These time and effort-intensive methods can only be applied to one disorder at a time. The different CDGs are likely connected because they all affect protein glycosylation and show similar symptoms in patients (65). Because of their relatedness, it is probable that common modifiers affect multiple CDGs. Here, we leveraged evolutionary tools to analyze all glycosylation genes and identify potential modifiers for all CDG genes. Further exploration of these common modifiers could lead to novel therapeutic targets for CDG patients that may be broadly applied to multiple types of CDGs.

## METHODS

### Evolutionary Rate Covariation

Evolutionary rate covariation (ERC) was calculated independently for orthologous sets of genes from: 62 mammal species, 39 non-mammalian vertebrate species, 22 drosophila species, 18 yeast species, and 17 nematode species. The gene-specific rate of evolution was calculated for each gene in all five independent phylogenies using the PAML package (66). The evolutionary rate for each gene was then normalized into relative evolutionary rates (RERs) by comparing the gene-specific branch length to the genome-wide average. The RERs for each pair of orthologous genes in each of the five datasets were correlated using a Pearson correlation. The correlation (i.e. the ERC value) was then Fisher transformed to adjust for the number of branches contributing to a score. This allowed us to sum all five datasets into one integrated FtERC dataset. The orthologs from drosophila, yeast, and nematodes were mapped to the hg19 human orthologs using InParanoid (67). Only 1:1 orthologs were retained, but we did not require that an ortholog be present in every independent phylogeny. This resulted in a dataset of FtERC values for 19,149 orthologous genes.

### Statistics

Significance of elevated ERC scores was determined using proteome-wide permutation testing. For significance testing in one group of proteins, the mean ERC value observed between all pairs in a group is calculated. Then it is compared to the mean ERC values of 100,000 groups with the same number of proteins chosen at random from the entire proteome. A p-value is calculated as the proportion of random groups of proteins that had a mean ERC value of equal or greater value than the mean of the tested group. Using 100,000 permutations, the most significant p-value possible is <1×10^-5^.

ERC significance between two different groups of proteins also used proteome-wide permutation testing. The mean ERC of scores between proteins in two different groups is calculated. Then it is compared to the mean ERC value between proteins where one of the testing groups is maintained, and the other group is substituted by the same number of proteins chosen at random from the entire genome. This is repeated 100,000 times, and a p-value is calculated as the proportion of random groups of proteins that had a mean ERC value of equal or greater value than the mean of the tested group. This process is repeated while maintaining the second group of proteins and replacing the first group with random proteins. The p-value for that testing is calculated as described above, and the higher p-value between the two tests is used.

Enrichment of each GPI anchor synthesis protein in the genome-wide dataset was determined using chi-squared testing to compare the number of observed occurrences of each protein in the dataset compared to number of expected occurrences if they were evenly distributed.

Genetic interactions between knockdown groups were determined using a two-way ANOVA run in RStudio. GPI anchor gene level (wild type or knockdown) and gene 2 gene level (wild type or knockdown) were the variables compared in the analysis.

### Gene Ontology Analyses

Gene Ontology (GO) analysis was performed using the GO enrichment analysis (geneontology.org) powered by the PANTHER Classification System. Enriched pathways were identified using the “GO biological process complete” annotation dataset, and enriched cell compartments were identified using the “GO cellular compartment complete” annotation dataset with the *Homo sapiens* gene reference list using the default parameters (Fisher’s Exact, Calculate False Discovery Rate). Figures 5 and 6 and S8 Fig show most specific subclass term as sorted by Panther hierarchical soring.

### Fly stocks and maintenance

Stocks were maintained on standard Archon Scientific glucose fly food at 25°C on a 12-h light/dark cycle. The *w-;; eya composite-GAL4* and *w-;eya composite-GAL4/CyO;+* lines were a gift from Justin Kumar (Indiana University Bloomington) and were characterized previously (38). *Drosophila* orthologues of candidate genes were determined using DIOPT, requiring a score ≥6 (68). RNAi lines used in this study were ordered from the Bloomington Drosophila Stock Center and the Vienna Drosophila Resource Center (Line IDs in S3 Data).

To create double knockdown flies, either *w-; GPI RNAi/CyO;eya-GAL4/TM3, Sb* or *w-;eya-GAL4/CyO; GPI RNAi/TM3, Sb* was crossed to either the candidate modifier RNAi lines or the equivalent control lines (*attP40, attP2, attP, w1118)* selecting progeny that were negative for both *CyO* and *TM3, Sb* balancers.

### Eye imaging and quantification

Adult female flies aged 3-5 days were collected under CO_2_ anesthesia then frozen at -80°C for later imaging. Eyes were imaged at 3x magnification using the Leica EC3 Camera. Eye area was measured as previously described (14,69).

### qPCR

Knockdown efficiency for each RNAi line used was tested using qPCR. RNAi lines were crossed to a *Tubulin-GAL4*/*Sb, Dfd-YFP* driver (BDSC: 5138) to test full-body knockdown amounts. 5-8 adult female flies aged 3-5 days old were collected, and RNA was extracted using a Direct-zol RNA Miniprep (Zymo Research R2061) using TRIzol Reagent (ThermoFisher Cat # 15596026) and including the DNAse step. RNA was converted to cDNA using a ProtoScript® II First Strand cDNA Synthesis Kit (NEB Cat # E6560L). RT-qPCR was performed using a QuantStudio 3 96-well 0.2 ml block instrument and PowerUp SYBR Green Master Mix (ThermoFisher Cat # A25741). If *Tubulin* driving the RNAi line was adult lethal, 5-10 YFP-negative larvae were used. If available, we used primers from the FlyPrimerBank (70) located at http://www.flyrnai.org/flyprimerbank. Other primers were designed using Primer3Plus (71) located at www.primer3plus.com. All primer sequences listed in S3 Data.

