## Supplementary figures and images for "Evolutionary rate covariation is pervasive between glycosylation pathways and points to potential disease modifiers"

### S1 Fig

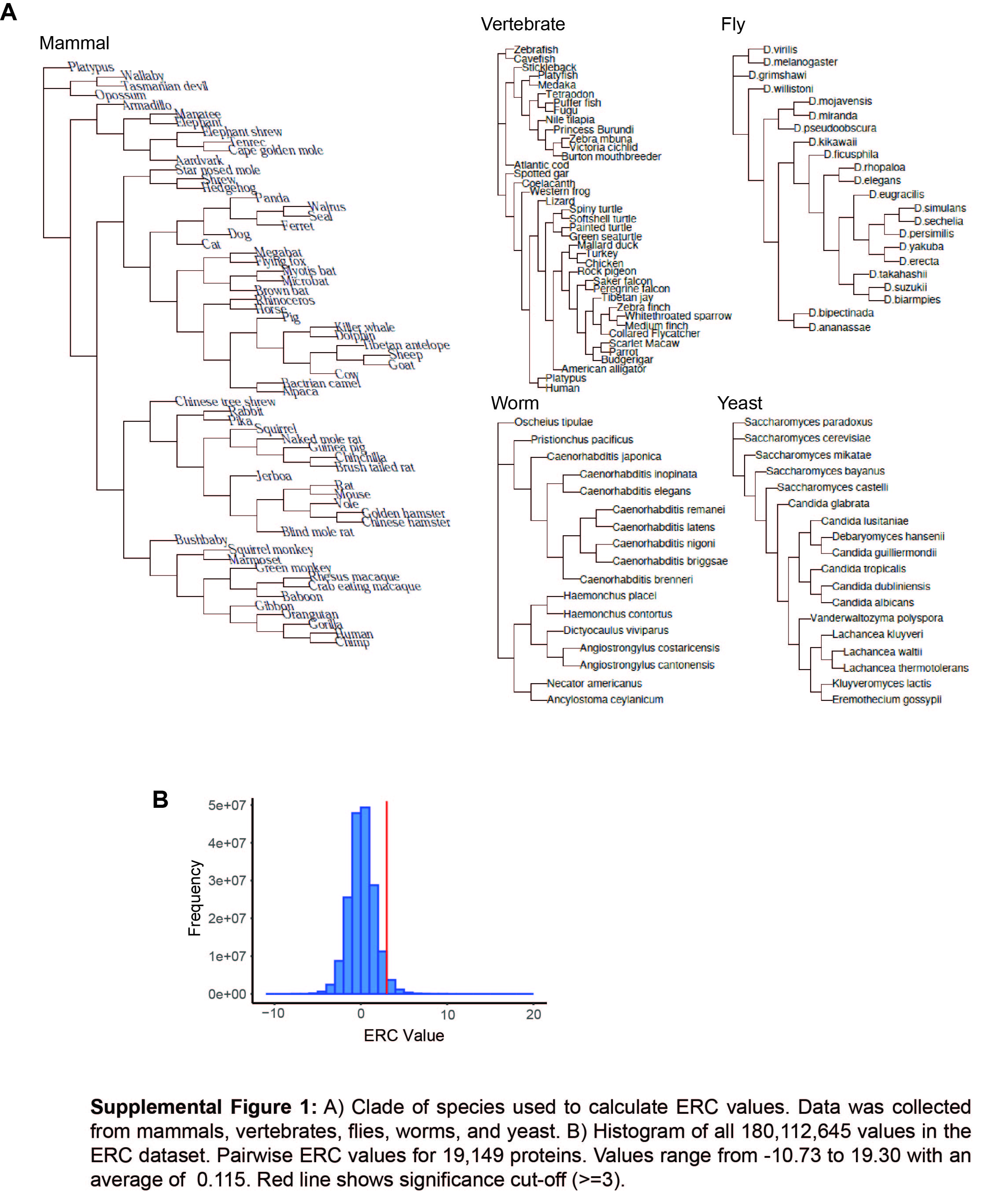

### S1 Table

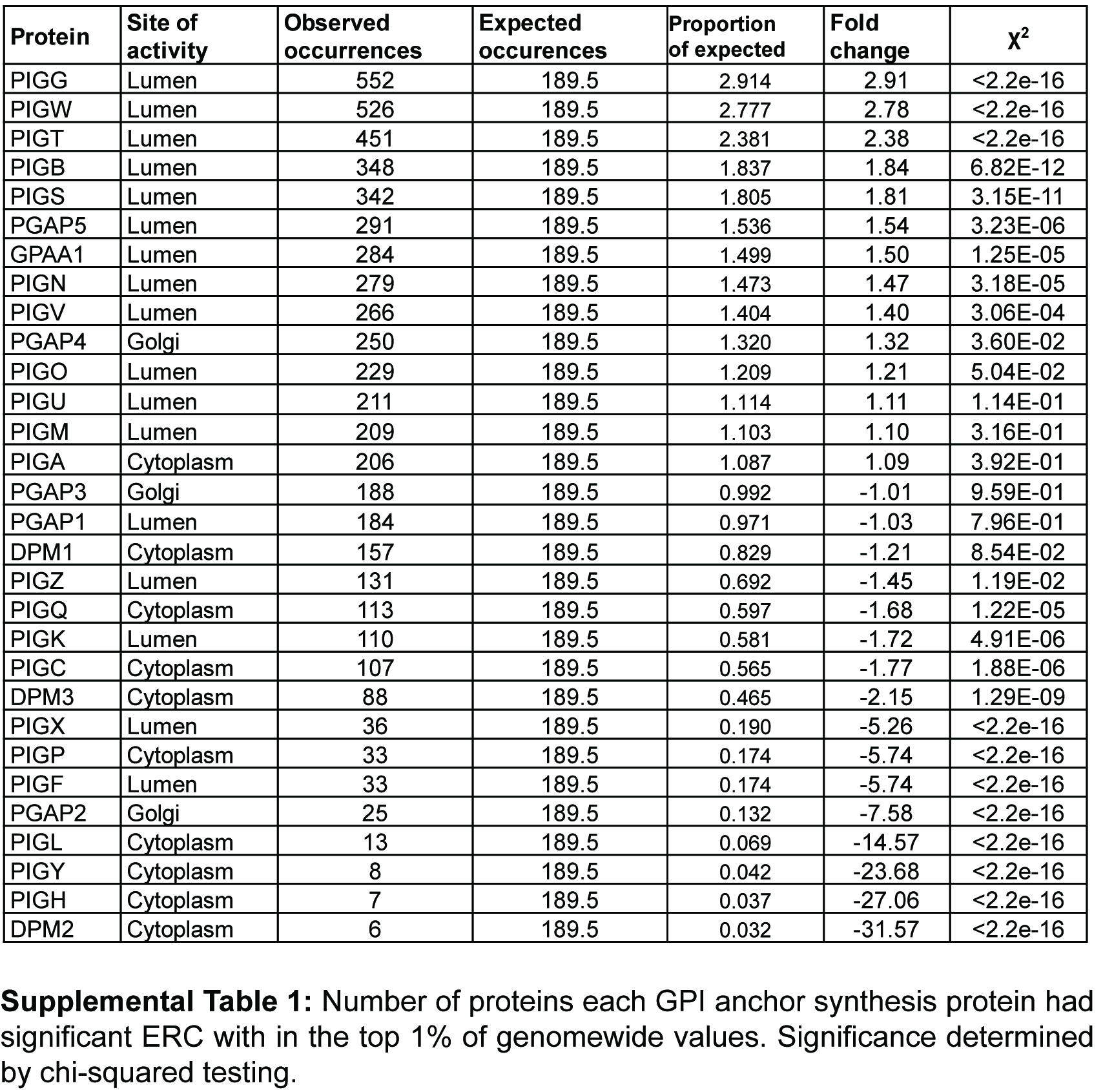

### S2 Fig

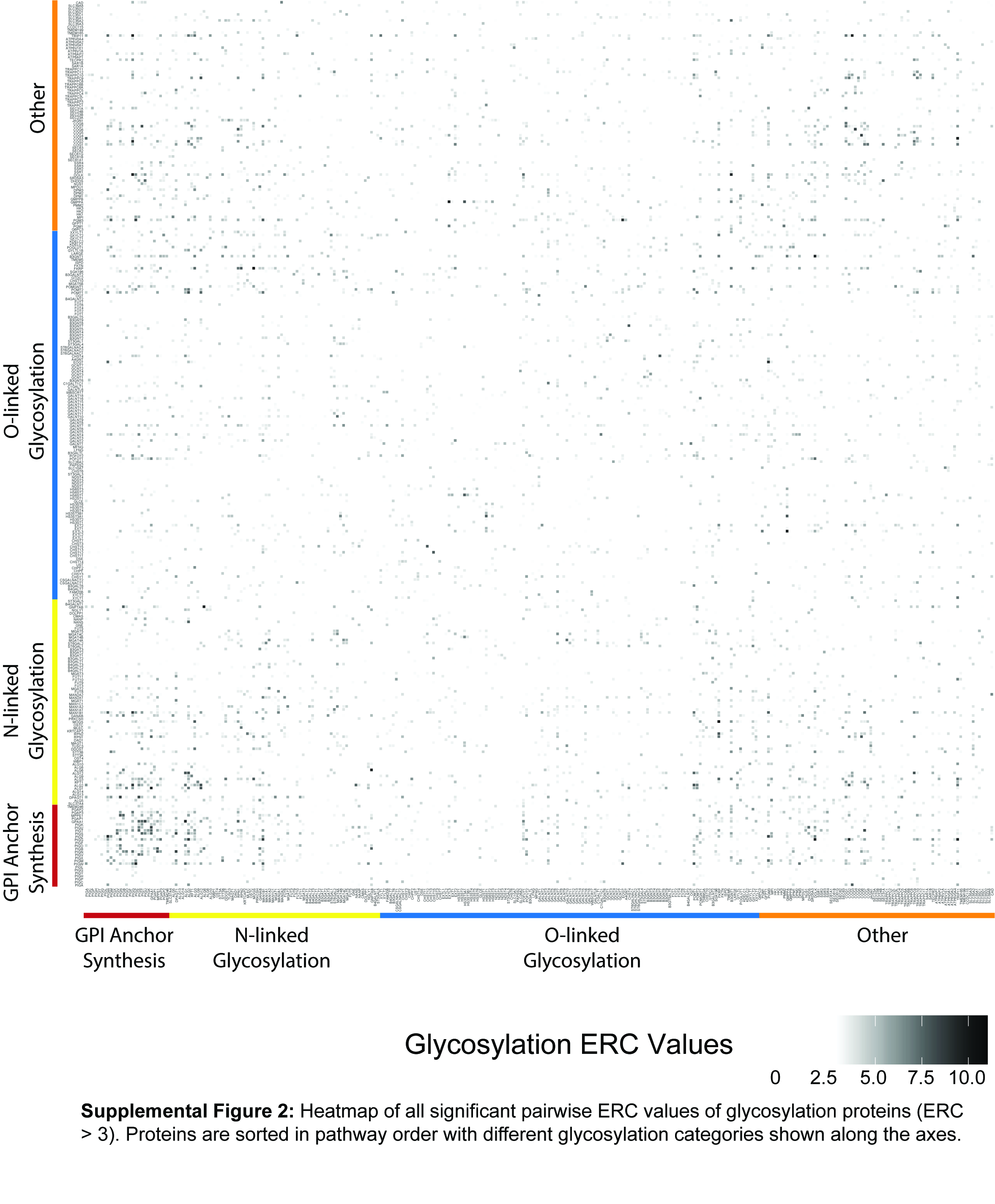

### S3 Fig

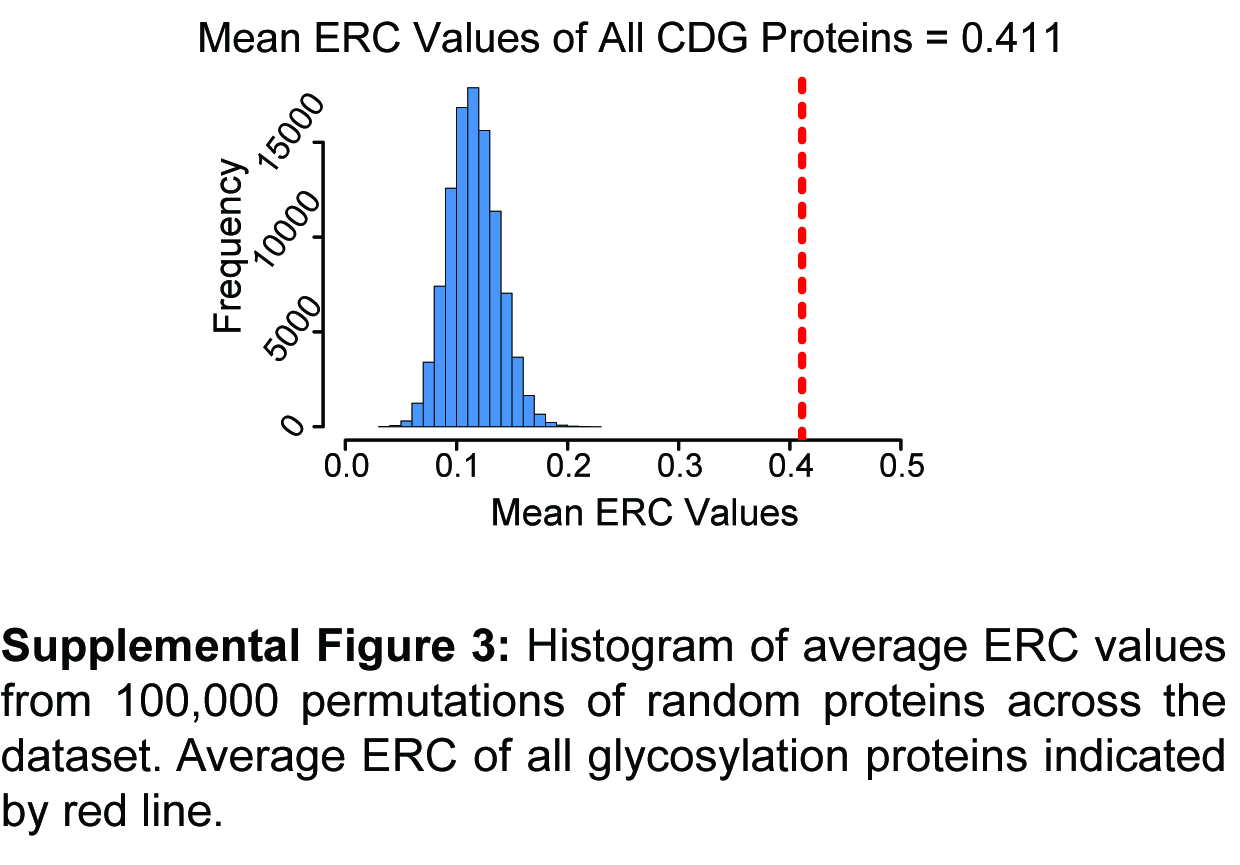

### S5 Fig

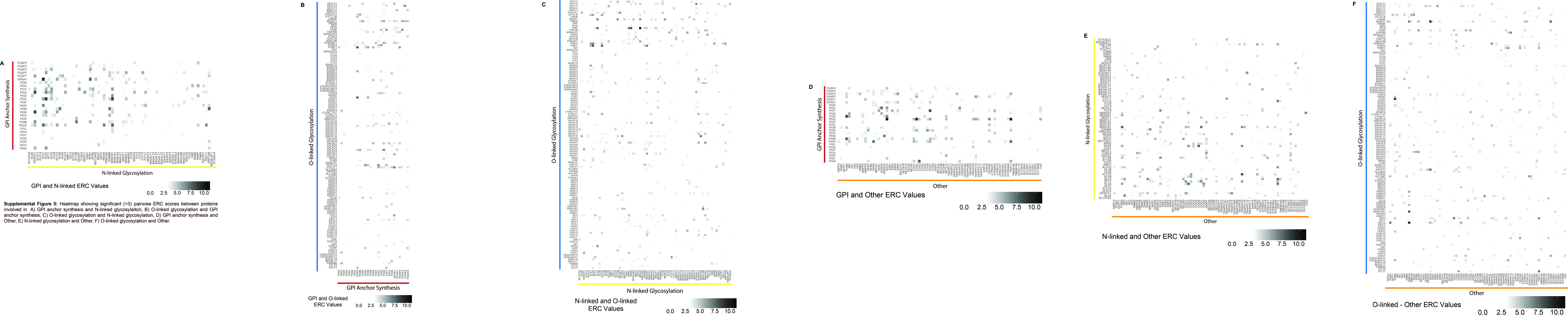

### S6 Fig

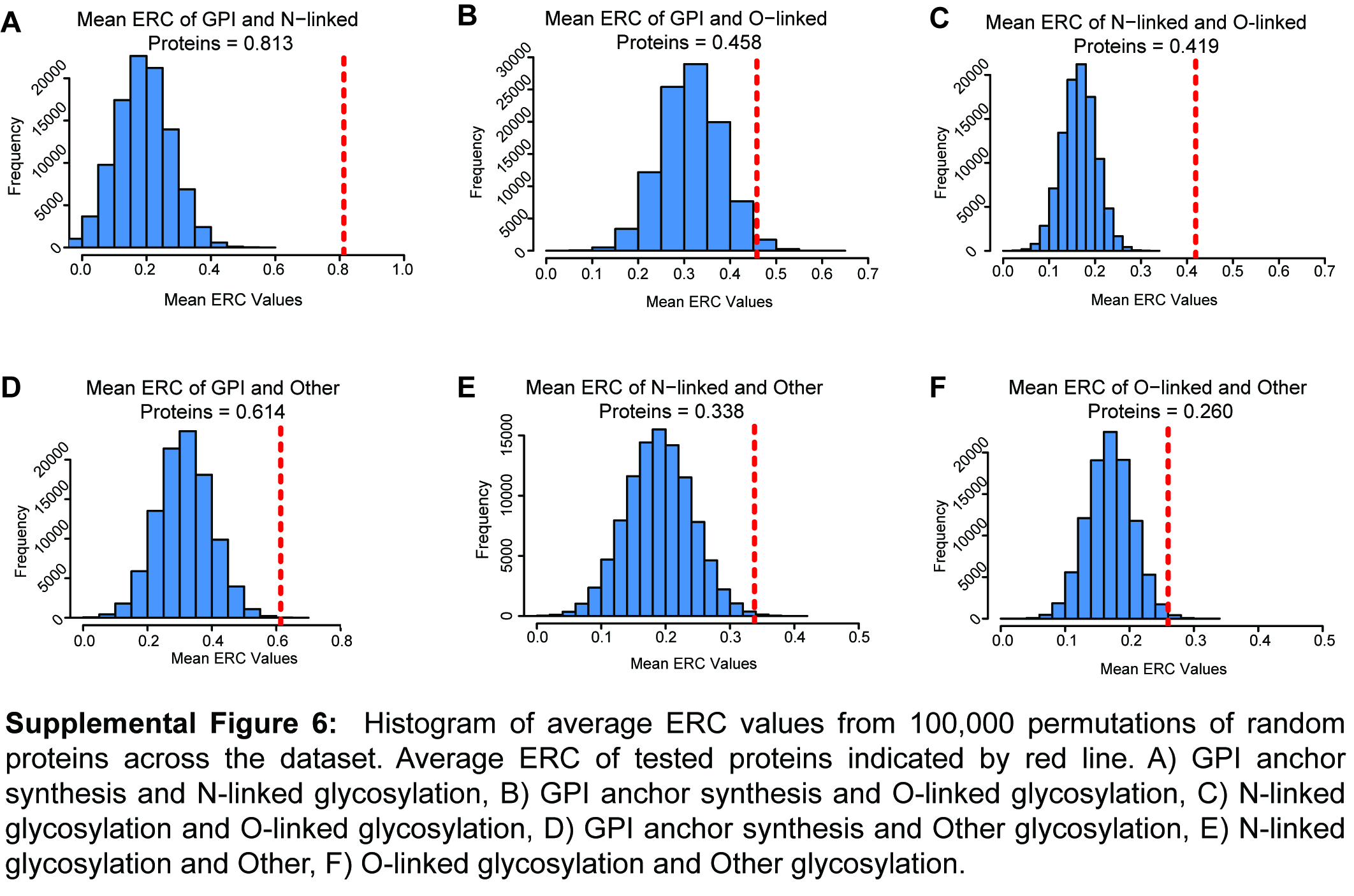

### S7 Fig

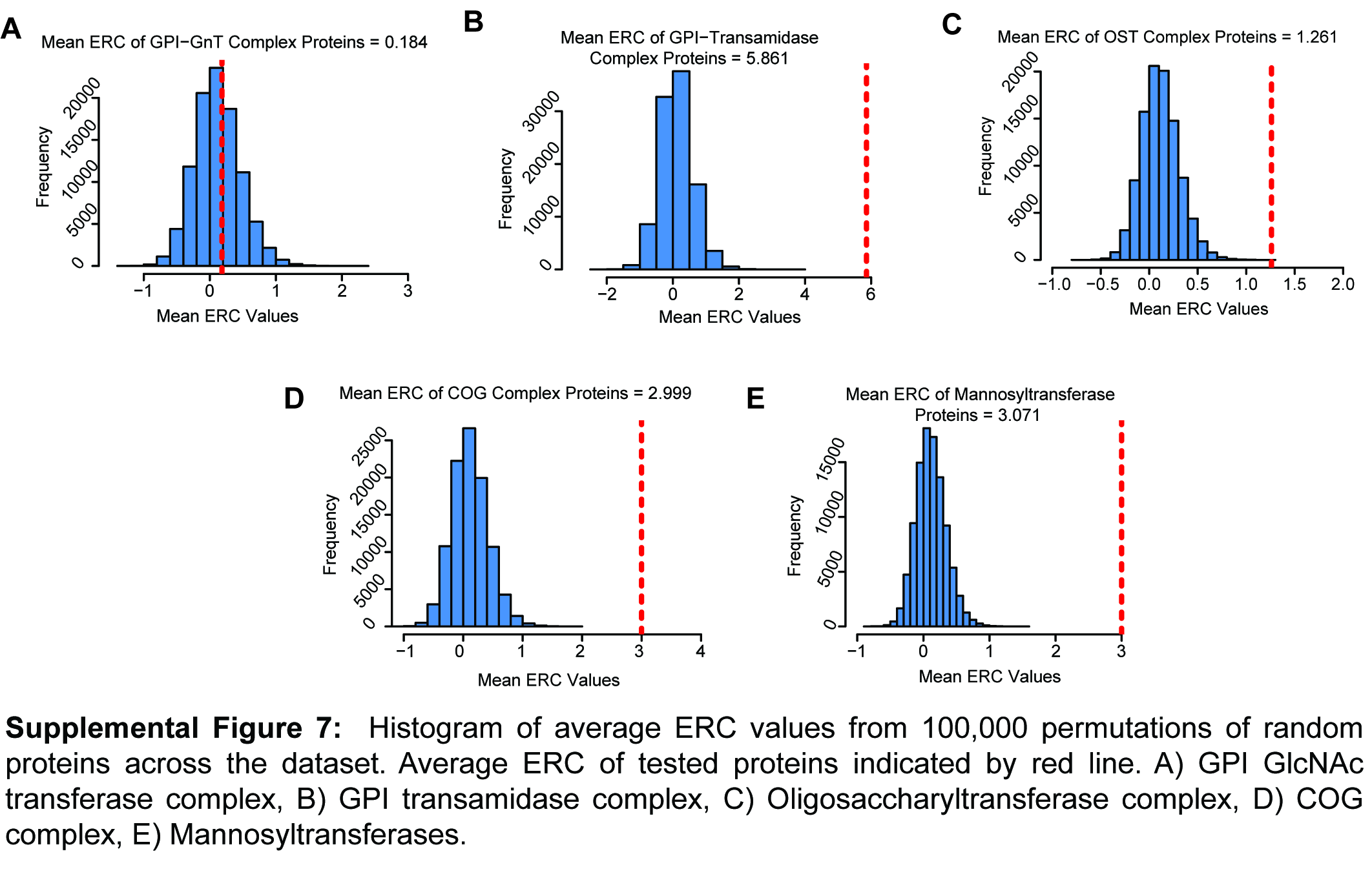

### S8 Fig

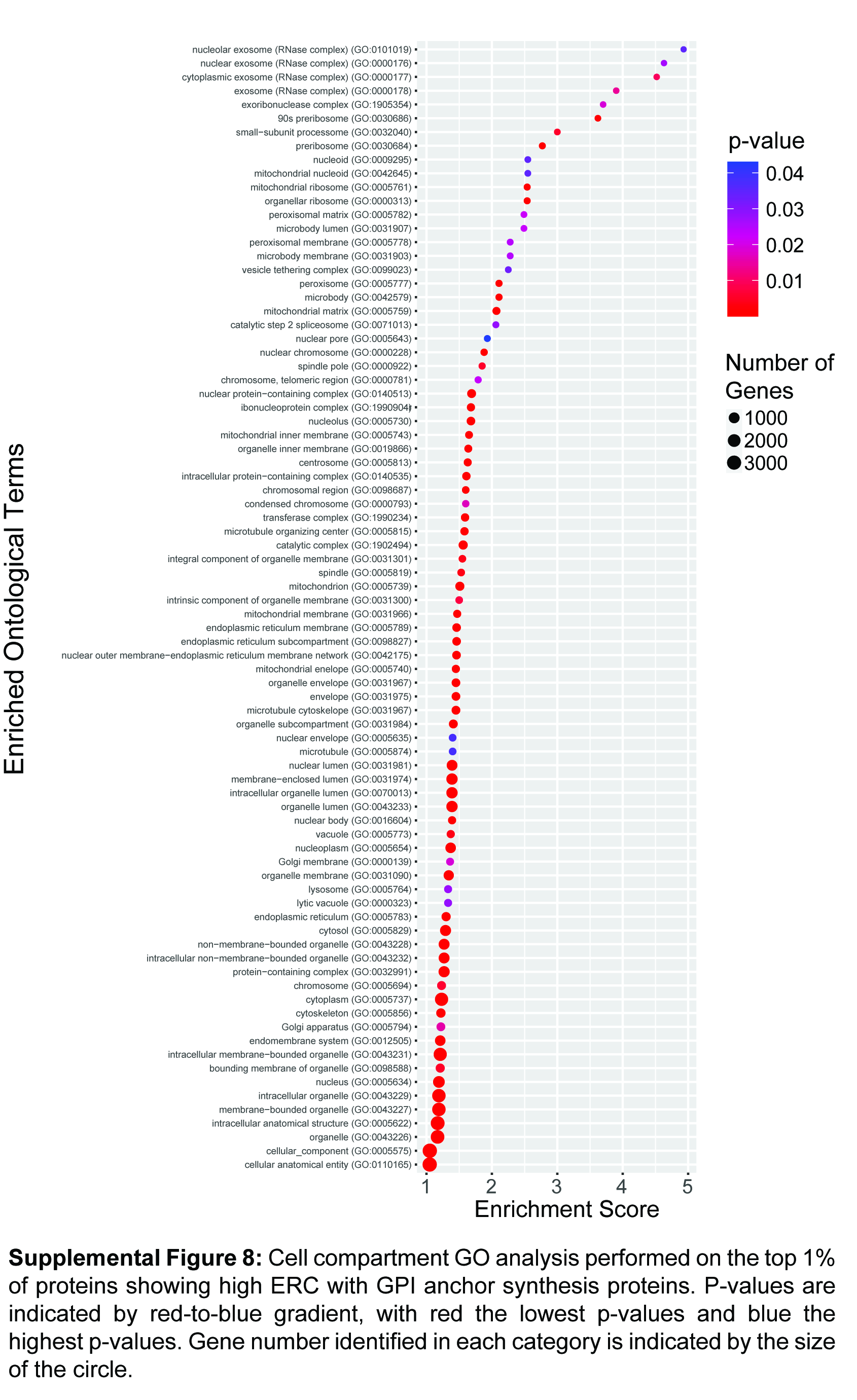

### S9 Fig

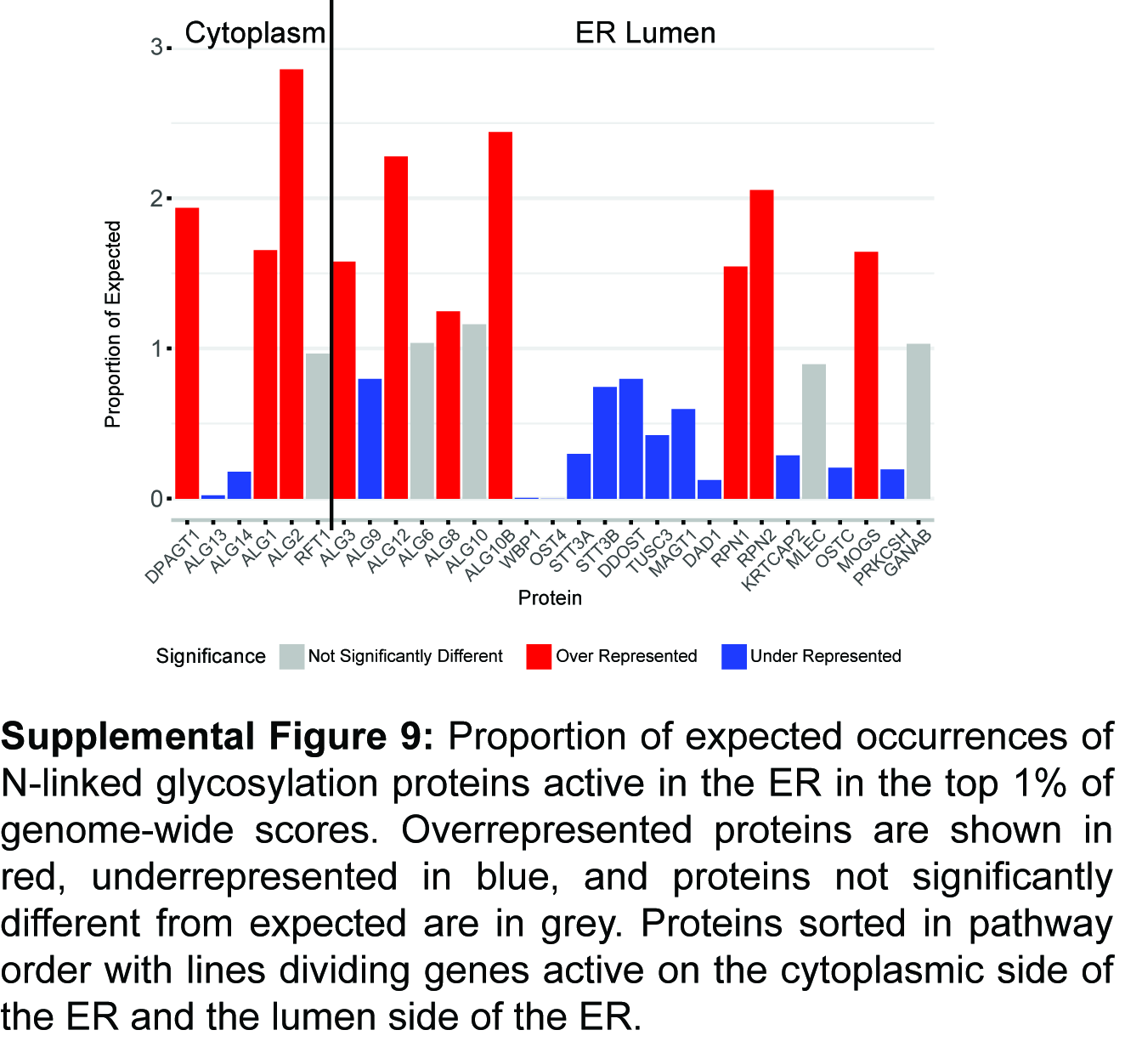
